# CO_2_ fixation mediated by the carbon concentrating mechanism enables a rapid response to nitrogen deprivation in cyanobacteria

**DOI:** 10.64898/2026.05.20.726413

**Authors:** Evan V. Saldivar, Julia Gershon, Juliana Artier, Dimitri Tolleter, Patrick M Shih, Seung Y. Rhee, Adrien Burlacot

## Abstract

Cyanobacteria are leading biomass producers of the ocean whose ecological success relies on their ability to respond to dynamic availability of nutrients like CO_2_ and nitrogen, which require distinct adaptive mechanisms. To survive nitrogen deprivation, cyanobacteria undergo a reversible transition to a dormant mode. Under low CO_2_ levels, a CO_2_ concentrating mechanism (CCM) supports their CO_2_ fixation. While the CCM and nitrogen assimilation have been shown to share some regulatory pathways, how the CCM impacts the response to nitrogen deprivation remains underexplored. In this study, by using mutants of the coastal cyanobacteria *Synechococcus* sp. PCC 7002 lacking a CCM component, we show that the high rate of carbon fixation mediated by the CCM tunes the speed of the nitrogen deprivation response in β-cyanobacteria. We first show that CCM mutants are deficient in inducing their typical nitrogen deprivation response under atmospheric CO_2_. However, at higher CO_2_ concentrations, the CCM mutants induce the nitrogen deprivation response. By combining Rubisco kinetics modeling with measurement of the response speed to nitrogen in various CO_2_ concentrations, we show that the speed of the nitrogen deprivation response increases linearly with Rubisco’s carboxylation rate. We further reveal that the regulation of nitrogen response by the CCM is also present in the distantly related freshwater cyanobacteria *Synechococcus elongatus* PCC 7942, suggesting a widespread role of this regulation across β-cyanobacteria. This study demonstrates that CO_2_ fixation by the cyanobacterial CCM is a key regulator of the nitrogen deprivation response, favoring a rapid response to dynamic environments.

## Introduction

Aquatic cyanobacteria are key primary producers responsible for an estimated 13-30% of global net primary production across marine and freshwater environments (Flombaum *et al*., 2013; Rae, Long, Badger, *et al*., 2013; Crockford *et al*., 2023). Their growth depends on the availability of nutrients, including carbon, phosphorus, and nitrogen, which is often limited in their native habitats (Droop, 1973; Schwarz and Forchhammer, 2005). Nutrient limitation events vary in timescale and can last on the order of hours and days, like nitrogen, which can be rapidly resupplied by the movement of nitrogen-rich water (Kubryakova and Korotaev, 2016), to millions of years, like CO_2_, which has steadily decreased over the last million years until the nineteenth century (Flamholz and Shih, 2020). Because each nutrient varies independently of the others in aquatic environments (Shaffer, 1989; Falkowski, Barber and Smetacek, 1998; Kubryakova and Korotaev, 2016; Sun *et al*., 2024), multiple modes of nutrient deprivation are possible, including deprivation of a single nutrient or simultaneous deprivation of multiple nutrients. Cyanobacteria have evolved to cope with these nutrient limitations to remain one of the most highly productive photoautotrophic clades.

How cyanobacteria respond to single-nutrient limitations is well-characterized. Depletion of major nutrients such as nitrogen (N), phosphorus (P), or sulfur (S) elicits similar responses in cyanobacteria (Allen and Smith, 1969; Ihlenfeldt and Gibson, 1975; Wanner *et al*., 1986; Collier and Grossman, 1992; Ariño *et al*., 1995). Although the responses to N, P, and S invoke distinct transcriptional programs (reviewed in Schwarz and Forchhammer (2005)), the three response pathways have several commonalities. One of the common responses is the degradation of phycobiliproteins (phycocyanin and allophycocyanin) that compose the light-harvesting complexes (phycobilisomes, PBS) (Allen and Smith, 1969). Termed chlorosis, this degradation of PBS (i) limits photosynthetic light energy capture and (ii) allows for the remobilization of a large internal pool of the nutrients sequestered in the PBS (Grossman *et al*., 1993). The rapid degradation of PBS is facilitated by the quick induction of small proteins, non-bleaching A and B (NblA and NblB), that destabilize the PBS complexes, permitting their proteolysis by proteases (Collier and Grossman, 1994) that degrade 50% of PBS between 6 and 12 hours post-deprivation for N and S, or P, respectively (Collier and Grossman, 1992). In contrast to the rapid proteolysis of PBS, the complete degradation of photosynthetic machinery, chlorophyll, and thylakoid membranes takes several days (Wanner *et al*., 1986; Collier and Grossman, 1992) and occurs after cells have accumulated carbon storage compounds, including poly-β-hydroxybutyrate (PHB) and glycogen granules (Klotz *et al*., 2016). These changes are reversible, and within days of N, P, or S resupply, cyanobacterial cells resume metabolism and growth by consuming the carbon stored as glycogen and PHB to fuel cellular respiration (Wanner *et al*., 1986; Klotz *et al*., 2016). Unlike the similarities in response to N, P, S limitation, cyanobacteria exhibit an entirely different nutrient deprivation response when CO_2_ decreases below the optimal levels for growth (1–3% CO_2_ in air) (Woodger, Badger and Price, 2005). In response to carbon limitation (<1% CO_2_), cyanobacteria maintain their PBS and chlorophyll, and use a carbon concentrating mechanism (CCM) to continue growing despite the nutrient deprivation.

The CCM achieves two main functions: (i) enhancing the CO_2_ fixation rate of Ribulose-1,5-bisphosphate carboxylase/oxygenase (Rubisco), and (ii) limiting Rubisco’s oxygenation reaction, which is energetically wasteful and results in net CO_2_ loss (Price and Badger, 1989, 1991; Badger and Price, 2003; Rae, Long, Badger, *et al*., 2013; Rae, Long, Whitehead, *et al*., 2013). These functions are achieved by two CCM components: a constitutive core and an inducible enhancement system that together enable efficient Rubisco function. One component of the constitutive CCM centers on the carboxysome, a proteinaceous microcompartment that encapsulates Rubisco and a Carbonic Anhydrase (CA) (Kerfeld and Melnicki, 2016). The CA functions to convert bicarbonate into CO_2_ and produces a high concentration of CO_2_ around the active site of Rubisco (Price and Badger, 1989). Because CA consumes bicarbonate within the carboxysome, the CA establishes a gradient of bicarbonate across the carboxysome shell that drives diffusion of inorganic carbon (C_i_) into the carboxysome. In this way, the carboxysome concentrates CO_2_ around Rubisco’s active site. When C_i_ limitation leads to a low concentration of bicarbonate in the cytoplasm, cyanobacteria actively transport bicarbonate across the cell membrane to supply the carboxysome with bicarbonate, using a set of constitutive and inducible C_i_ transporters, completing the CCM (Badger and Price, 2003; Hagemann, Song and Brouwer, 2021). The combined function of both CCM components is essential for cellular growth at ambient CO_2_; the genetic ablation of either the carboxysome or membrane C_i_ transporters is sufficient to disrupt the CCM (Badger and Price, 2003). Most cyanobacterial samples from natural environments are proposed to have an active CCM (Badger and Price, 2003). Thus, most current nutrient deprivation events in nature occur in cyanobacterial cells with an active CCM. However, little is known about how the function of the CCM impacts other nutrient deprivation responses.

The CCM has been proposed to contribute to signaling required for the N-deprivation response (Forchhammer and Selim, 2020). However, evidence supporting this hypothesis is limited. It was early recognized that N deprivation-mediated PBS degradation is inhibited in the absence of inorganic carbon (Wood and Haselkorn, 1980), suggesting that C_i_ availability impacts N deprivation-mediated PBS degradation. And more recently, mutant cells lacking the carboxysome have been shown to have a partially-inhibited PBS degradation phenotype during nitrogen starvation (Luo *et al*., 2025). However, this partial PBS degradation phenotype in a CCM mutant was observed under high CO_2_ conditions where the CCM is not necessary (Luo, 2025). This study therefore probes the relationship between the CCM and PBS degradation during N deprivation at a CO_2_ condition where the CCM is not required for C_i_ fixation. Indeed, while it has been proposed that the CCM could have an effect on N-deprivation response pathways by either (i) increasing CO_2_ fixation rates (Forchhammer and Selim, 2020; Luo *et al*., 2025) or (ii) limiting the oxygenation reaction of Rubisco (Woodger, Badger and Price, 2005), evidence supporting either hypothesis is missing.

Here, by measuring the PBS content of mutants of the CCM at various CO_2_ levels, we show that CCM function is required for the N-deprivation response. By varying CO_2_ and O_2_ levels during N deprivation, we further establish that the carbon fixation rate of Rubisco, not the oxygenation rate, correlates with the speed of PBS degradation during N deprivation. This study establishes the CCM as a key regulator of the N-deprivation response, which, when active, ensures rapid PBS degradation across a broad range of CO_2_ availability.

## Results

### The CCM is required for nitrogen deprivation-mediated phycobilisome degradation in β-cyanobacteria

As the CCM’s role is more apparent at ambient CO_2_ (0.04% CO_2_; **LC, Table 1**), we hypothesized that its role under N deprivation (**Fig. 1A**) would be most apparent under LC. We thus grew *Synechococcus sp*. PCC 7002 mutants harboring a deletion of the CCM operon required for carboxysome formation (referred to here as Δ*ccm-1*, previously generated in Abernathy *et al*., (2019); Hill *et al*., (2020), **Table 2**), and transitioned them from N-replete media at high CO_2_ (3% CO_2_; **HC, Table 1**) to N deprivation at LC (**Fig. 1B-E**). Following 24 hours of N deprivation, wild-type (**WT-1**) cells became chlorotic, while little visible change could be seen in Δ*ccm-1* cells (**Fig. 1B**). The change in Δ*ccm-1* cells after 24 hours was indistinguishable between N-deplete and N-replete conditions (**Sup Fig. S1A, B**). In line with this result, the absorbance spectra of whole cells at wavelengths corresponding to PBS (620-650 nm) (Apt, Collier and Grossman, 1995) were substantially reduced in WT-1 compared to Δ*ccm-1* cells after 24 hours of N deprivation (**Fig. 1C**). The absorbance was similar in WT-1 and Δ*ccm-1* cells before N deprivation (**Sup Fig. S1C**). In addition, in the presence of N, no difference was observed in absorbance between the WT-1 and Δ*ccm-1* cells after 24 hours at LC (**Sup Fig. S1**), suggesting that the decrease of absorbance in WT-1 was specific to N deprivation. WT-1 showed a 75% reduction in PBS content per cell after N deprivation at LC, while Δ*ccm-1* cells did not significantly change (**Fig. 1D, E**). To confirm that the CCM ablation caused the absence of bleaching in N deprivation for Δ*ccm-1*, we conducted the same experiment with an independently-generated CCM mutant (**Δ*ccm-2***) in WT-2 background (**Table 2**) as well as a CCM mutant harboring a CCM operon under the control of an Isopropyl-β-D-galactopyranoside-(IPTG) inducible promoter (**CCMi**, Hill *et al*., (2020), **Table 2**). After 24 hours of N deprivation in LC, both WT-2 and CCMi induced with IPTG showed strong PBS degradation (**Sup Fig. S2**). However, Δ*ccm-2* and the non-IPTG-induced CCMi did not show a PBS content decrease. Taken together, we conclude that the PBS decrease characteristic of N deprivation depends on the presence of a functional CCM at LC.

**Table 1.** Gas mixtures used throughout this study. Calculations for bicarbonate concentrations in the media are detailed in (**Methods**). Bicarbonate concentrations assume equilibrium and Per Rubisco carboxylation rate and oxygenation rate were calculated using the Rubisco kinetic model (**Methods**).

| Name | Gas Composition | Estimated Bicarbonate Concentration (mM) | Carboxylation Rate (Rubisco<sup>-1</sup> /sec<sup>-1</sup>) | Oxygenation Rate (Rubisco<sup>-1</sup>/sec<sup>-1</sup>) |
| --- | --- | --- | --- | --- |
| LC | 0.04% CO<sub>2</sub>,<br>21% O<sub>2</sub>, 79% N<sub>2</sub> | 0.4 | 0.44 | 0.14 |
| Intermediate 1 | 0.27% CO<sub>2</sub>,<br>21% O<sub>2</sub>, 79% N<sub>2</sub> | 3 | 2.5 | 0.11 |
| MC | 0.7% CO<sub>2</sub>,<br>21% O<sub>2</sub>,<br>79% N<sub>2</sub> | 8 | 5.0 | 0.09 |
| Intermediate 2 | 1.4% CO<sub>2</sub>,<br>21% O<sub>2</sub>, 78% N<sub>2</sub> | 15 | 7.13 | 0.07 |
| HC | 2.5% CO<sub>2</sub>,<br>21% O<sub>2</sub>, 76.5% N<sub>2</sub> | 30 | 9.6 | 0.04 |

**Table 2.** Description and design of strains in this study.

| Name | Genetic Background | Modification to Background | Reference |
| --- | --- | --- | --- |
| WT-1 | *Synechococcus* sp. PCC 7002 (PCC 7002) | None | Pasteur Collection of Cyanobacteria |
| $\Delta ccm-1$ | WT-1 | Ablation of Carbon Concentrating Mechanism -carboxysome operon (*ccm*) locus | (Abernathy *et al.* 2019) (original name $\Delta ccm$) |
| WT-2 | WT-1 | Replacement of Rubisco Large Subunit (RbcL) replaced with RbcL-GFP | (Hill *et al.* 2020) (original name RbcL-GFP) |
| $\Delta ccm-2$ | WT-2 | Ablation of *ccm* locus in RbcL-GFP background ($\Delta ccm$-RbcL-GFP) | (Hill *et al.* 2020) (original name $\Delta ccm$) |
| CCMi | $\Delta ccm-2$ ($\Delta ccm$-RbcL-GFP) | Addition of *ccm* locus under Isopropyl-β-D-galactopyranoside (IPTG) inducible promoter | (Hill *et al.* 2020) (original name $\Delta ccm^+$) |
| WT-7942 | *Synechococcus elongatus* PCC 7942 (PCC 7942) | None | Pasteur Collection of Cyanobacteria |
| $\Delta ccm$-7942 | WT-7942 | Ablation of Carbon Concentrating Mechanism -carboxysome operon (*ccm*) locus | This study |

**Figure 1.**
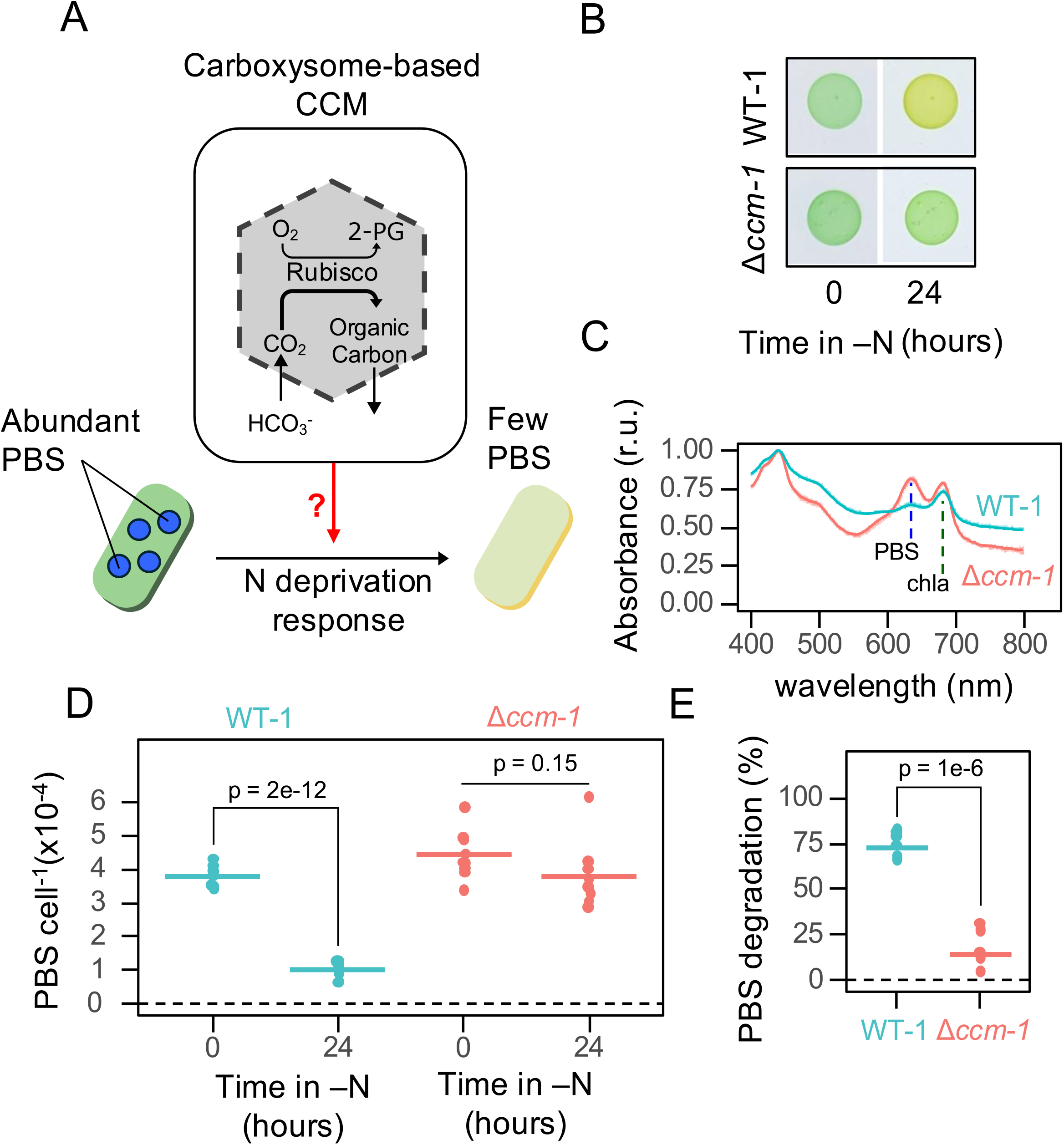
The CCM is required for PBS degradation after N deprivation. A) Schematic of the cyanobacterial CCM (top) with a focus on carbon fixation by Rubisco. A schematic of the N-deprivation response (bottom) with a representative vegetative cell (left) transitioning to a dormant state (right) following N deprivation. The central question, whether the CCM regulates the N-deprivation response, is depicted with a red arrow. Acronyms: carbon dioxide (CO_2_), bicarbonate (HCO_3_^-^), carbon concentrating mechanism (CCM), phycobilisomes (PBS), nitrogen (N), 2-phosphoglycolate (2-PG) B) Representative images of WT-1 and Δ*ccm-1* cells taken before and after 24 hours of N deprivation on solid medium at 0.04% CO_2_. C) Absorbance spectra for WT-1 and Δ*ccm-1* cells following 24 hours of N deprivation in liquid culture at 0.04% CO_2_. Absorbance values were rescaled so that max absorbance = 1. Two vertical dashed lines show representative wavelengths used to estimate the abundance of PBS and chlorophyll A (chla). Lines represent means of 3 independent biological replicates. Shown are absorbance spectra representative of three biological replicates. D) Quantified PBS content for WT-1 and Δ*ccm-1* cells following 0 and 24 hours of N deprivation in liquid culture at 0.04% CO_2_. PBS content was normalized by cell count. E) The relative decrease in per-cell PBS content (PBS degradation) calculated from values in D. In (D) and (E), each dot represents a single biological replicate, and horizontal bars represent the mean. Data shown are from 3 independent experiments, with 2–3 biological replicates per genotype per experiment (*N = 3, n = 2-3*). p-values from a Student’s t-test are shown.

### Rubisco’s carboxylase reaction, not oxygenase activity, tunes PBS degradation in Δ*CCM*

We hypothesized that the non-bleaching phenotype of Δ*ccm-1* could be complemented by increasing C_i_ availability, which rescues the growth of CCM mutant cells in N-replete media (Price and Badger, 1989; Hill *et al*., 2020) (**Sup Fig. S3**). To test this, we measured PBS degradation following 24 hours of N deprivation at CO_2_ levels ranging from LC (0.04% CO_2_) to high CO_2_ (2.5% CO_2_ in air, referred to as **HC**). In WT-1, PBS degradation was indistinguishable across all tested CO_2_ conditions, with a mean PBS degradation of 75% (**Fig. 2**). However, PBS degradation was dependent on CO_2_ availability in the Δ*ccm-1* mutant, influencing both the level (**Fig. 2**) and speed (**Fig. 3**) of PBS degradation. Negative PBS degradation values, observed in Δ*ccm-1* at LC (**Fig. 3**), could reflect minor changes in PBS levels or cell density at LC (**Sup Fig. S3**). Most notably, WT-1 cells showed indistinguishable PBS degradation dynamics at LC and HC, while Δ*ccm-1* cells demonstrated negligible PBS degradation at LC, but comparable PBS degradation to WT-1 at HC (**Fig. 3**). Although this result demonstrates a relationship between CO_2_ availability and the speed of PBS degradation in Δ*ccm-1*, a mechanistic explanation remains undemonstrated.

**Figure 2.**
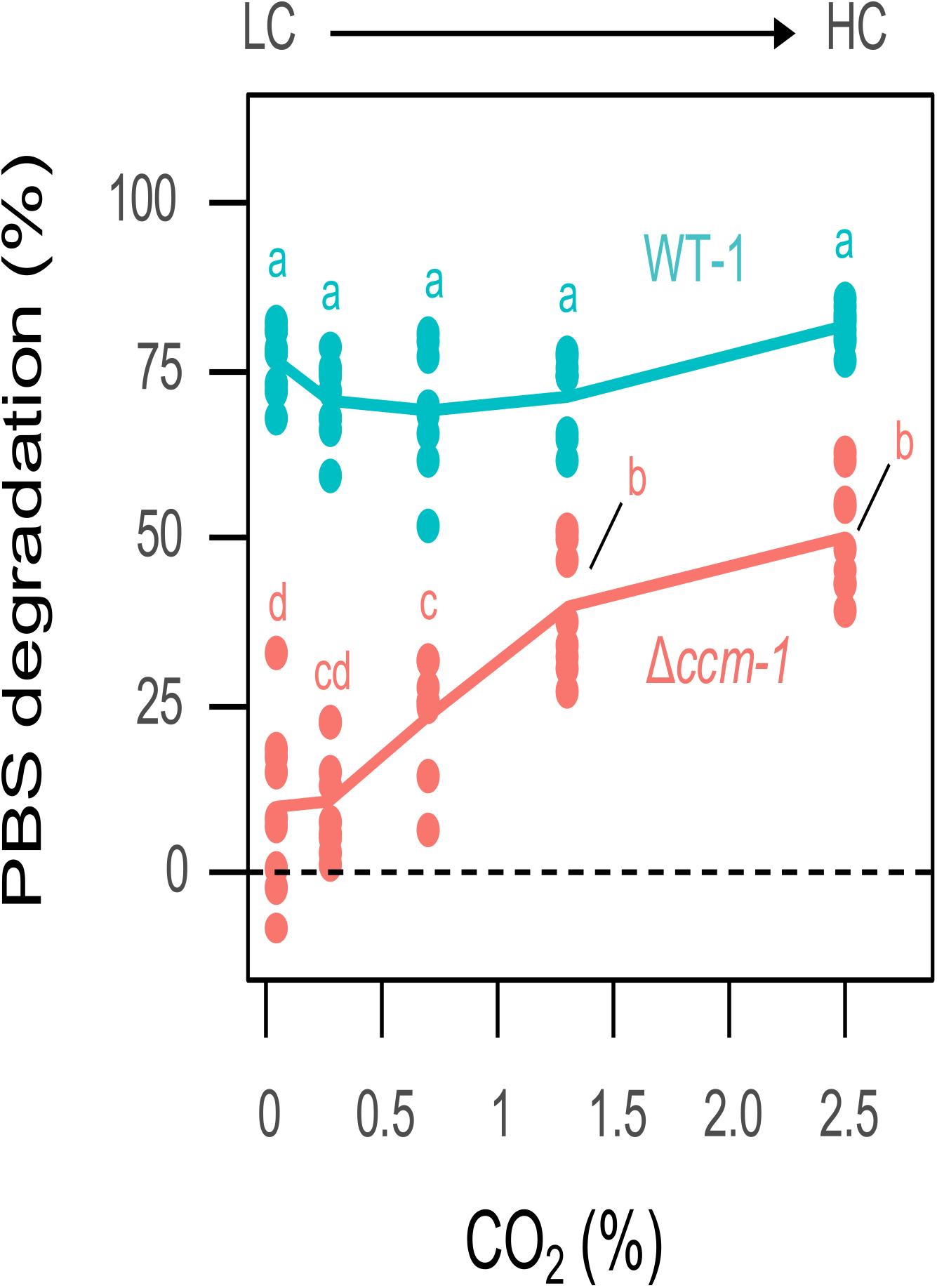
N deprivation-mediated PBS degradation in CCM mutant cells increases with CO_2_ concentration. PBS degradation after 24 hours of N deprivation for WT-1 and Δ*ccm-1* cyanobacteria across various CO _2_ conditions. Tested CO_2_ concentrations are 0.04% (LC), 0.23%, 0.7%, 1.3%, and 2.5% (HC). Data are from *N = 3* independent experiments with (*n = 3* biological replicates per genotype per condition per experiment (*N = 3, n = 3*). Each data point represents a single biological replicate. The line indicates the mean of all biological replicates. Letters represent significantly different groups at p-value < 0.05 from a two-way ANOVA and Tukey’s Honest Significance Difference (HSD) test.

**Figure 3.**
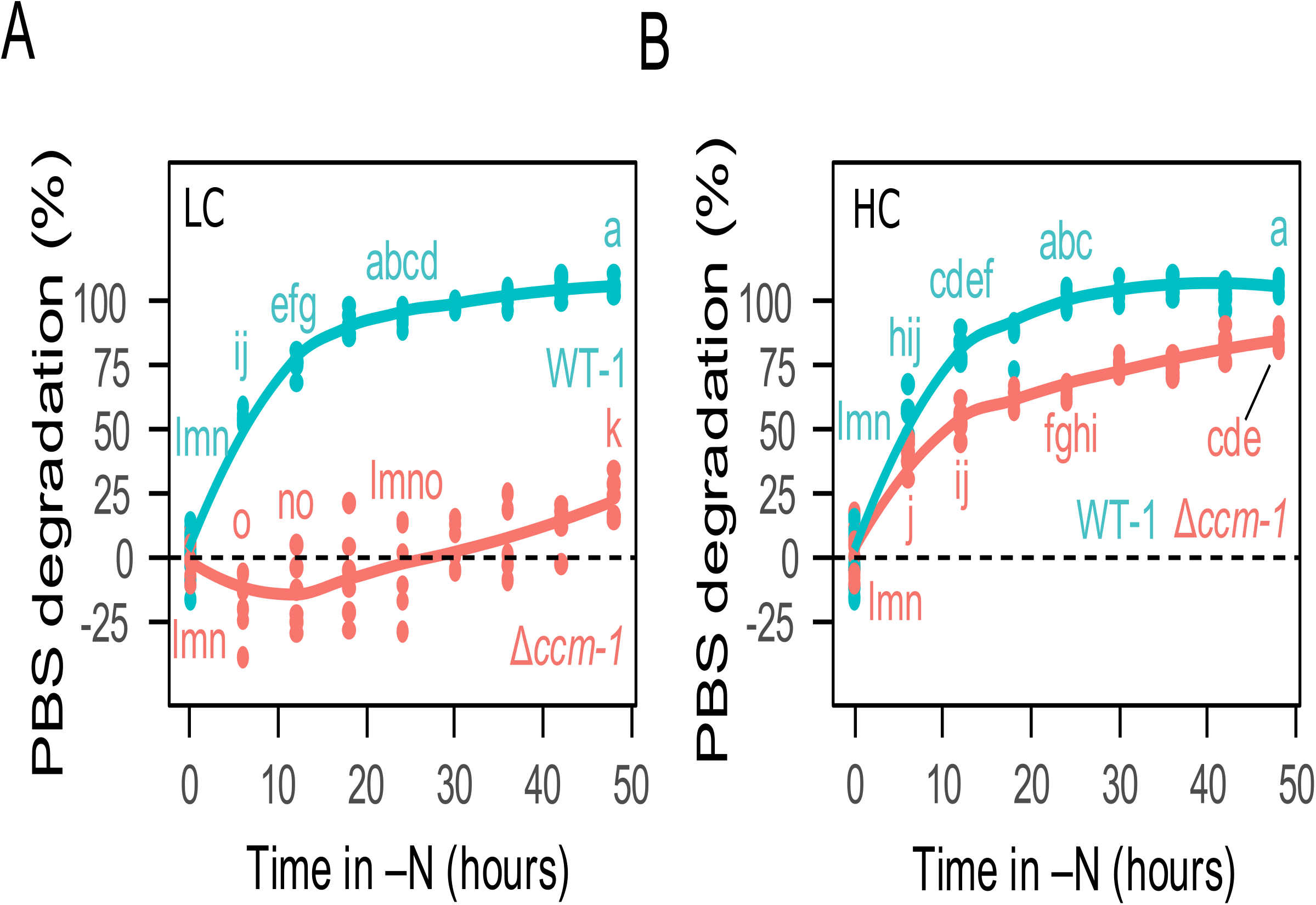
CO_2_ availability modulates the dynamics of PBS degradation in CCM mutants. Dynamics of PBS degradation during nitrogen (N) deprivation at 0.04% CO_2_ (LC, A) and 3% CO_2_ (HC, B). Each data point represents a single biological replicate. Solid lines indicate the mean of all biological replicates from *N = 2* independent experiments each with *n = 3* biological replicates per genotype. Letters represent significance codes from a three-way ANOVA and Tukey’s Honest Significance Difference (HSD) test performed across (A) and (B). Different letters indicate a statistically significant difference at p-value < 0.05.

Adding CO_2_ results in the accumulation of C_i_ in the media as both CO_2_ and bicarbonate. We next sought to understand which form of C_i_ causes PBS degradation in Δ*ccm-1*. To test if bicarbonate causes PBS degradation in Δ*ccm-1*, we added up to 30 mM sodium bicarbonate, which is the estimated amount of bicarbonate in HC media (**Table 1**), to N-deplete media at LC. Δ*ccm-1* did not degrade PBS in response to bicarbonate addition (**Sup Fig. S4**), suggesting that CO_2_, not bicarbonate, drives PBS degradation in Δ*ccm-1*. Because CCM mutant cells fix CO_2_ but not bicarbonate (Price and Badger, 1989; Hill *et al*., 2020), we hypothesized that carbon fixation drives PBS degradation in Δ*ccm-1*. To test this, we measured PBS degradation in the absence of light at HC. PBS degradation significantly decreased for both Δ*ccm-1* and WT-1 cells in the dark, as compared to being in the light (**Sup Fig. S5**), suggesting that the light energy required for CO_2_ assimilation is necessary for N-deprivation mediated PBS degradation in both strains. We therefore hypothesized that the CO_2_-dependent carbon fixation drives PBS degradation in β-cyanobacteria in response to N deprivation.

To characterize the relationship between CO_2_ fixation and PBS degradation, we used established kinetic parameters for Rubisco to model carboxylation rates (**Fig. 4A, B**) across the CO_2_ conditions used in **Fig. 2**. We found a significant linear relationship between the predicted Rubisco-mediated carboxylation rate and PBS degradation in Δ*ccm-1* (**R**^**2**^ **= 0.71, p-value = 3.38 x 10**^**-13**^, **Fig. 4C, Table 3**). As a control, we modeled the activity of the second major cyanobacterial carboxylase, phosphoenolpyruvate carboxylase (**PEPC**), across the CO_2_ conditions used in this study (**Sup Fig. S6**). The linear relationship between PEPC activity and PBS degradation was much weaker (**R**^**2**^ **= 0.53, Sup Fig. S6, Table 3**) than for Rubisco. We therefore conclude that PBS degradation is most likely driven by Rubisco activity and not PEPC in the conditions tested in this study.

**Table 3.** Comparison of PBS degradation to Rubisco and PEPC activity. Modeled relationships comparing N deprivation-mediated PBS degradation to Rubisco carboxylation, Rubisco oxygenation, and PEPC carboxylation rates. Estimate (β) denotes the linear model’s estimate of the slope. The t-value corresponds to the test statistic for the null hypothesis that β = 0, and the p-value indicates the statistical significance of this test.

| Enzyme (Reaction) | Estimate ($\beta$) | Standard Error | t value | p value | Adjusted R<sup>2</sup> |
| --- | --- | --- | --- | --- | --- |
| Rubisco (carboxylation) | 5.057 | 0.486 | 10.40 | 3.38e-13 | 0.714 |
| Rubisco (oxygenation) | -486.309 | 50.777 | -9.58 | 4.04e-12 | 0.6785 |
| PEPC (carboxylation) | 50.527 | 8.952 | 5.64 | 1.29e-06 | 0.4178 |

**Figure 4.**
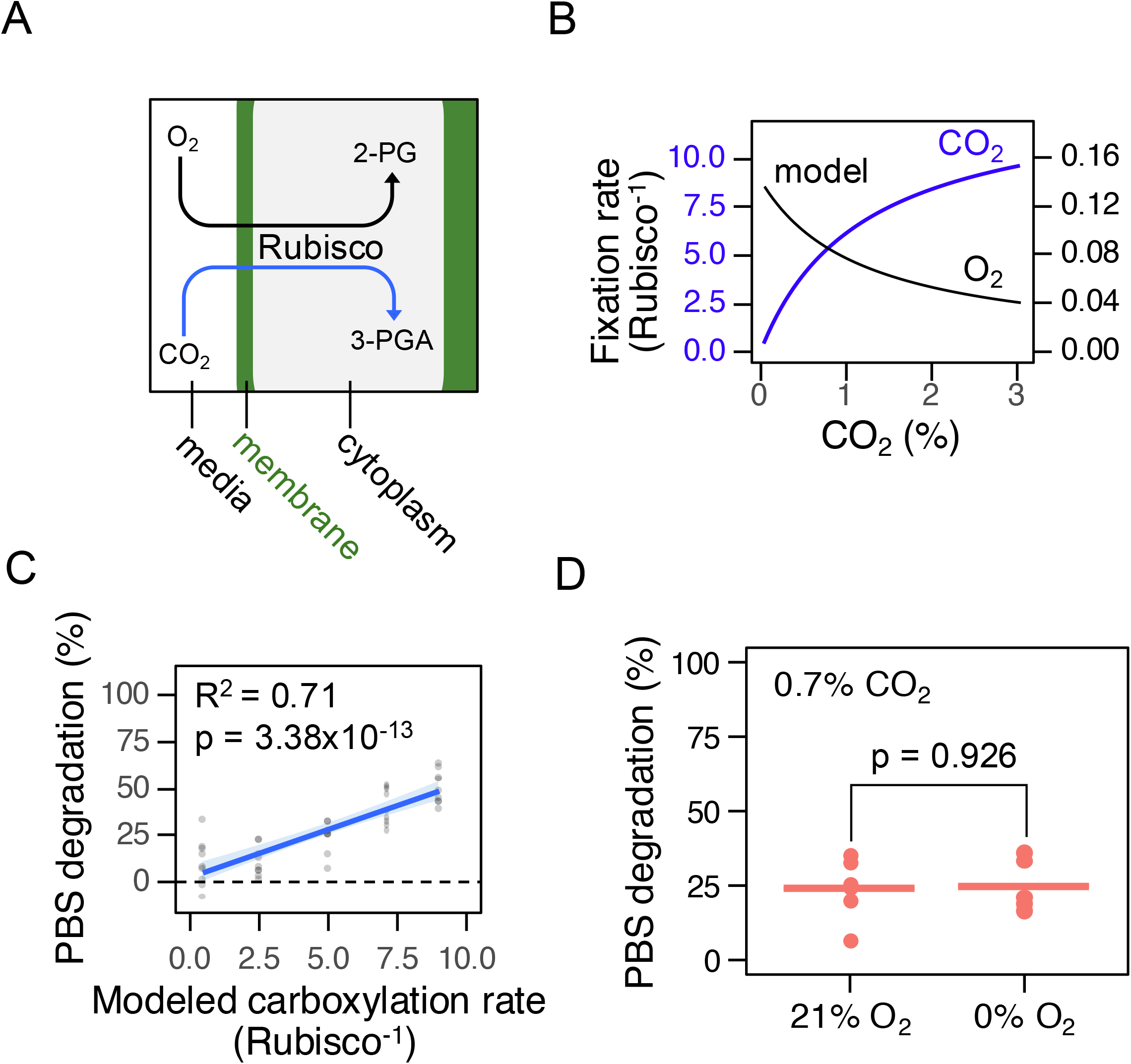
Rubisco carboxylase activity correlates with N deprivation-mediated PBS degradation in CCM mutants. A) Graphical description of the model for Rubisco in Δ*ccm-1* cells. The two compartments (media and cytoplasm) are separated by membranes (green border) that allows free diffusion of CO_2_ and O_2_. Acronyms: 2-PG, 2-phosphoglycolate; 3-PGA, 3-phosphoglycerate. B) Modeled carboxylation (blue) and oxygenation (black) rates per Rubisco calculated using kinetic parameters specific to PCC 7002 Rubisco (Cummins et al. 2018). C) Comparison of PBS degradation in Δ*ccm-1* cells (data from **Fig. 2**) with predicted Rubisco carboxylation rate from panel B. A best-fit linear regression (blue line) is shown, along with Adjusted R^2^ and the significance (p) of the F-statistic. The grey shading represents the 95% confidence interval. Data from **Fig. 2** are overlaid in light grey. D) PBS degradation after 24 hours of N deprivation at 0.7% CO_2_, with and without oxygen. Each data point represents a single biological replicate (*n = 3* biological replicates per genotype per condition per experiment, *N = 2* experiments). The horizontal line represents the mean of 9 biological replicates. The p-value from Student’s t-test is shown (p). The data in panel D were corrected to remove experimental batch effects. Raw data for panel D are provided in **Sup Fig. S7**.

While Rubisco carboxylase activity correlates linearly with PBS degradation in Δ*ccm-1* over the CO_2_ conditions tested, Rubisco oxygenase activity decreases as a function of CO_2_ availability (**Table 3**). Furthermore, the linear fit between Rubisco oxygenase activity and PBS degradation is as strong as the relationship between Rubisco carboxylase activity and PBS degradation (**Table 3)**. We therefore sought next to test if Rubisco oxygenase activity affects PBS degradation in Δ*ccm-1*. To accomplish this, we compared PBS degradation at 0.7% CO_2_ mixed with air (+ O_2_) and N_2_ gas (-O_2_) (**Fig. 4D**). PBS degradation was indistinguishable for Δ*ccm-1* with and without oxygen at 0.7% CO_2_, suggesting that Rubisco oxygenation does not impact the PBS degradation phenotype in the conditions tested (**Fig. 4D**).

### The role of the CCM in tuning the speed of N-deprivation response is also observed in a distantly-related *Synechococcus* species

We next sought to understand if the relationship between CCM activity and PBS degradation is specific to PCC 7002. To test if the relationship between PBS degradation and CO_2_ level is found in other cyanobacteria, we repeated the experiment in an additional CCM mutant we developed in the freshwater strain *Synechococcus elongatus* PCC 7942 (**Δ*ccm-7942*, Table 2**). Consistent with the coastal strain *Synechococcus* sp. PCC 7002, between LC and HC, PBS degradation increased from 0% to 75% in Δ*ccm-7942* (**Fig. 5**). At HC, both Δ*ccm-7942* and the WT (**WT-7942**) degraded 75% of their PBS content after 24 hours of N deprivation (**Fig. 5**). Based on these results, we conclude that the CO_2_-dependent PBS degradation occurs in CCM mutant strains of two distinct *Synechococcus* species, suggesting that the CCM-dependent regulation of N deprivation-induced PBS degradation may be conserved across β-cyanobacteria.

**Figure 5.**
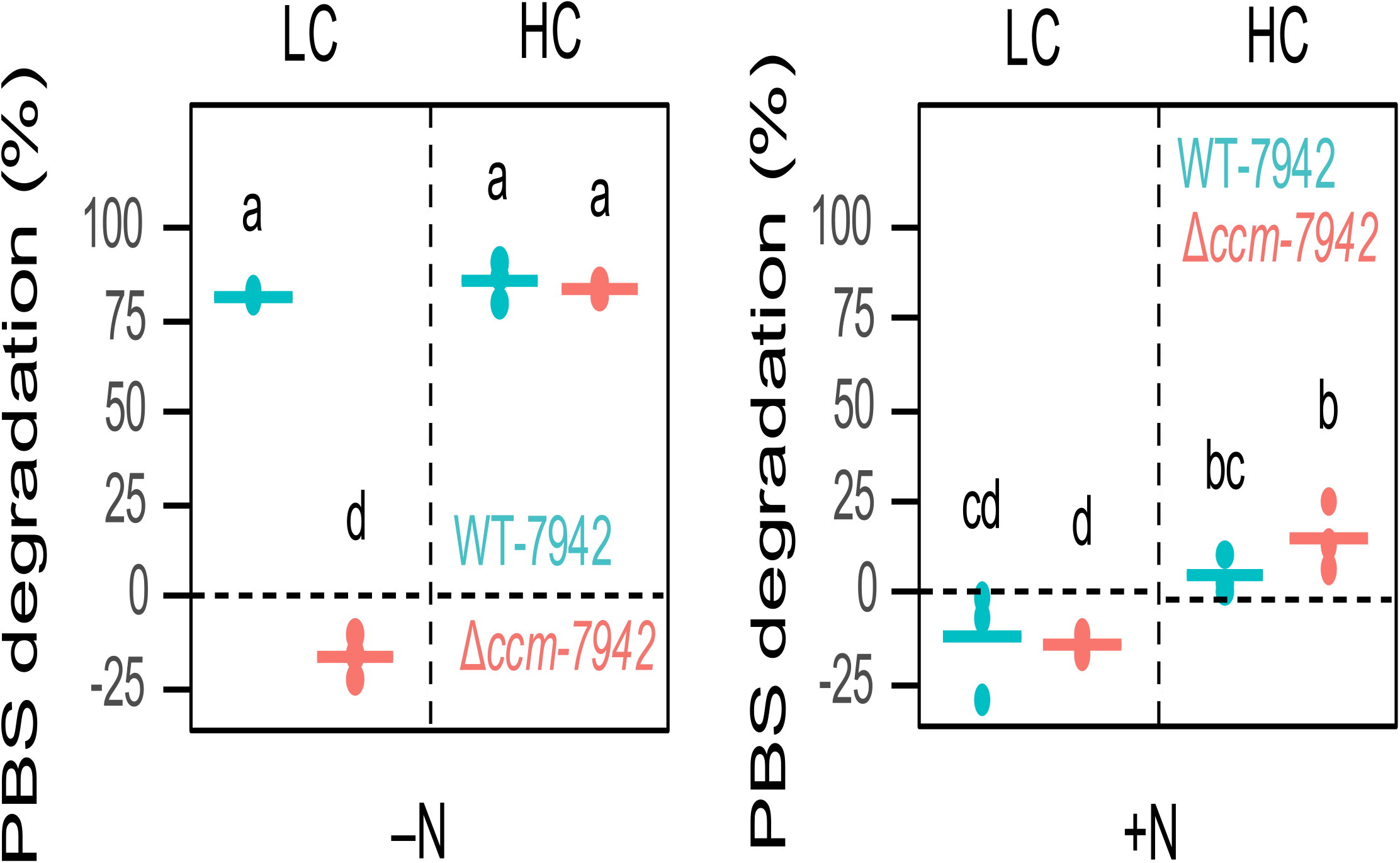
CO_2_ dependent PBS degradation in Synechococcus elongatus PCC 7942 (PCC 7942). PBS degradation after 24 hours in (N)-deplete media (left) and N-replete media (right) at 0.04% CO_2_ (LC) and 3% CO_2_ (HC) in WT-7942 and Δ*ccm-7942* strains in the PCC 7942 genetic background. Each data point represents a single biological replicate from one representative experiment (*n = 3* biological replicates per genotype per condition). The horizontal line indicates the mean of the three biological replicates. Letters represent significance codes from a three-way ANOVA and Tukey’s Honest Significance Difference (HSD) test. Different letters indicate a statistically significant difference at p-value < 0.05.

## Discussion

Previous studies have hypothesized that the CCM regulates the onset of N deprivation-mediated PBS degradation through CO_2_ fixation (Luo *et al*., 2025) or by limiting O_2_ generation (Forchhammer and Selim, 2020). Our study establishes that the CCM regulates N deprivation-mediated PBS degradation mostly through controlling the flux of CO_2_ fixed by Rubisco (**Figs. 1, 2, 4**). CCM mutants have been reported to exhibit limited PBS degradation during N deprivation (Luo *et al*., 2025). However, the investigation of CCM mutant behavior from Luo *et al*., (2025) was based on a single high CO_2_ level where CCM function is not required for cell growth. Furthermore, we show that the decreased PBS degradation of Δ*ccm-1* compared to WT-1 at HC (**Fig. 2**) is specific to PCC 7002, and was not reproduced in PCC 7942 (**Fig. 5**). Thus, our study expands on the findings of Luo *et al*., (2025) to show that CCM function most directly impacts N deprivation-mediated PBS degradation at low CO_2_ conditions, where the CCM is most relevant for growth, across multiple species. Additionally, Luo *et al*., (2025) showed that bicarbonate strongly influences N deprivation-mediated PBS degradation. While our findings suggest that bicarbonate does not influence PBS degradation (**Sup Fig. S4**), these data do not contradict Luo *et al*., 2025. Luo *et al*., (2025) showed that bicarbonate rescues PBS degradation in WT cells in the complete absence of C_i_, whereas our findings demonstrate that bicarbonate addition does not rescue PBS degradation at LC (**0.04% CO**_**2**,_ **0.4 mM bicarbonate, Table 1**) in Δ*ccm-1* cells. Thus, the two studies test the relationship between bicarbonate availability and N deprivation-mediated PBS degradation under different contexts that prevent direct comparison. Our findings reveal that multiple CCM mutants exhibit a strong CO_2_-dependent PBS degradation during N deprivation, and that Rubisco’s carboxylation, rather than PEPC’s carboxylation, predicts PBS degradation, expanding our understanding of this phenomenon. Beyond this, we show that Δ*ccm-1* cells degrade PBS at a rate determined by CO_2_ availability (**Fig. 3**) through a light-dependent process (**Fig. S5**), consistent with a Rubisco-dependent mechanism (**Fig. 4**). Further, we note that Rubisco’s oxygenation reaction does not regulate PBS degradation in the conditions tested. Because WT-1 cells do not change PBS degradation in response to CO_2_ (**Fig. 2**), we conclude that the CCM ensures a robust N-deprivation response in WT cells. To better understand the relationship between CCM function and the N-deprivation response, experiments probing different CCM components, such as C_i_ transporters, will be an essential path forward. Through this study of a single CCM component, we conclude that carbon fixation, enabled by carboxysome function, is a requirement for the N deprivation-mediated PBS degradation in WT-1.

In this study, the relationship between PBS degradation and CO_2_ availability for CCM mutants was observed in two distant β-cyanobacteria strains in the genus *Synechococcus*. Intriguingly, the conservation of this phenomenon across two distantly related *Synechococcus* species suggests that the CCM may be important for N deprivation-induced PBS degradation broadly across β-cyanobacteria. It has been noted that all extant cyanobacteria genetically encode components of the CCM (Axen, Erbilgin and Kerfeld, 2014; Sutter *et al*., 2021). This suggests that there is a strong selective pressure to maintain the CCM in many natural environments. However, the CCM is not strictly required in all environments – aquatic systems with elevated CO_2_ levels, such as near hydrothermal vents (Shitashima, 1998), would permit the growth of cyanobacteria lacking a CCM. It is therefore unclear why all reported cyanobacteria encode a CCM locus (Axen, Erbilgin and Kerfeld, 2014; Sutter *et al*., 2021). The relationship between the N-deprivation response and the CCM shown in this study, particularly in PCC 7002, suggests that even in the presence of sufficient CO_2_, acclimation to N deprivation occurs more slowly in cyanobacteria lacking a CCM compared to the WT counterparts (**Fig. 3**). This complex interaction between CCM function, CO_2_ fixation, and the N-deprivation response could thus explain part of the selective pressures that maintain the CCM in nature.

The results from this study also show that the CCM plays an essential role in the response to combined C_i_ and N deprivation in the environment. This is a relevant scenario in the environment, as both C_i_ and N are predicted to become depleted in environments where cyanobacterial productivity is high, such as during oceanic algal blooms (Engel *et al*., 2002). Although algal blooms are often caused by the addition of nitrogen to marine environments, active nitrogen uptake by algae rapidly drives environmental nitrogen rapidly back down to low levels (Engel *et al*., 2002). Thus, the initial stages of an algal bloom represent high N availability and strong C_i_ assimilation by the CCM, whereas the late stages represent deprivation of both C_i_ and nitrogen. During an algal bloom, the CCM therefore plays two roles: permitting rapid growth when nitrogen becomes available and inducing PBS degradation during nitrogen deprivation. The extent to which the interplay between C_i_ and N deprivation-mediated PBS degradation mediated by the CCM is required for the fitness of blooming cyanobacteria remains to be studied.

PBS degradation has been hypothesized as a requirement to survive N deprivation (Klotz and Forchhammer, 2017). In support of this hypothesis, most cyanobacterial mutants lacking the ability to degrade PBS are unable to survive N deprivation (Gründel *et al*., 2012; Klotz and Forchhammer, 2017; Díaz-Troya *et al*., 2020). The requirement for PBS degradation to survive N deprivation hinges on cellular energetics: N deprivation creates an imbalance between reductants produced by photosynthesis and used by N assimilation pathways (Collier and Grossman, 1992, 1994). Cyanobacterial cells therefore require downregulation of photosynthetic energy production during N deprivation to avoid cellular damage by reductants. Despite this general trend, we note that Δ*ccm-1* can survive N deprivation for up to 48 hours (**Sup Fig. S8**). This is surprising, as both C_i_ and N deprivation represent physiological states in which reductants generated by photosynthesis cannot be effectively dissipated. Further study is required to understand how cyanobacterial cells lacking a CCM withstand the pressures of photosynthetic light energy and electron flow in the absence of both nitrogen and carbon assimilation.

In this study we show that Rubisco and CCM function are essential components of the N-deprivation response in cyanobacteria. This suggests that, in some photosynthetic organisms, Rubisco has a role in tuning nutrient stress responses. This must be taken into account when engineering Rubisco to improve plant productivity: for crop engineering efforts to improve CO_2_ fixation abilities, such as adding CCMs or improving the specificity or speed of Rubisco (Slattery and Ort, 2014; South and Cavanagh, 2018), it will be important to assess their potential implications for resilience to nutrient changes.

## Limitations of this study

We observe that CCM mutant cyanobacteria do not completely degrade their phycobilisomes under high CO_2_ conditions (**Fig. 2**); however, this phenotype is specific to PCC 7002 and could not be recapitulated in PCC 7942 (**Fig. 5**). Thus, the primary strain in which we report a Rubisco carboxylation–dependent PBS degradation phenotype exhibits both a CO_2_-dependent and a CO_2_-independent component of PBS degradation. We do not currently understand the basis of the CO_2_-independent component in PCC 7002, but note that it is highly strain-specific, unlike the CO_2_-dependent PBS degradation.

We note that this study assumes that the gas concentrations around the Rubisco active site match the gas concentrations present in the media. However, myriad biological processes produce and consume O_2_ and CO_2_, and therefore, the effective intracellular concentrations of these gases may differ from the external environment. In particular, the production of O_2_ through photosynthesis could drive an increased intracellular O_2_ concentration. Without a method to measure the concentrations of O_2_ and CO_2_ in the immediate proximity of Rubisco, these modeled values are assumed to reflect conditions *in vivo*.

## Materials and methods

### Strains and growth conditions

Strains WT-1 and Δ*ccm-1* were developed and described previously in (Abernathy et al. 2019) (**Table 2**). Strains WT-2, Δ*ccm-2*, and CCMi were developed and described in Hill et al. (2020). Genotypes were confirmed using primers and PCR conditions from Hill et al. (2020) (**Table 4**).

**Table 4.**
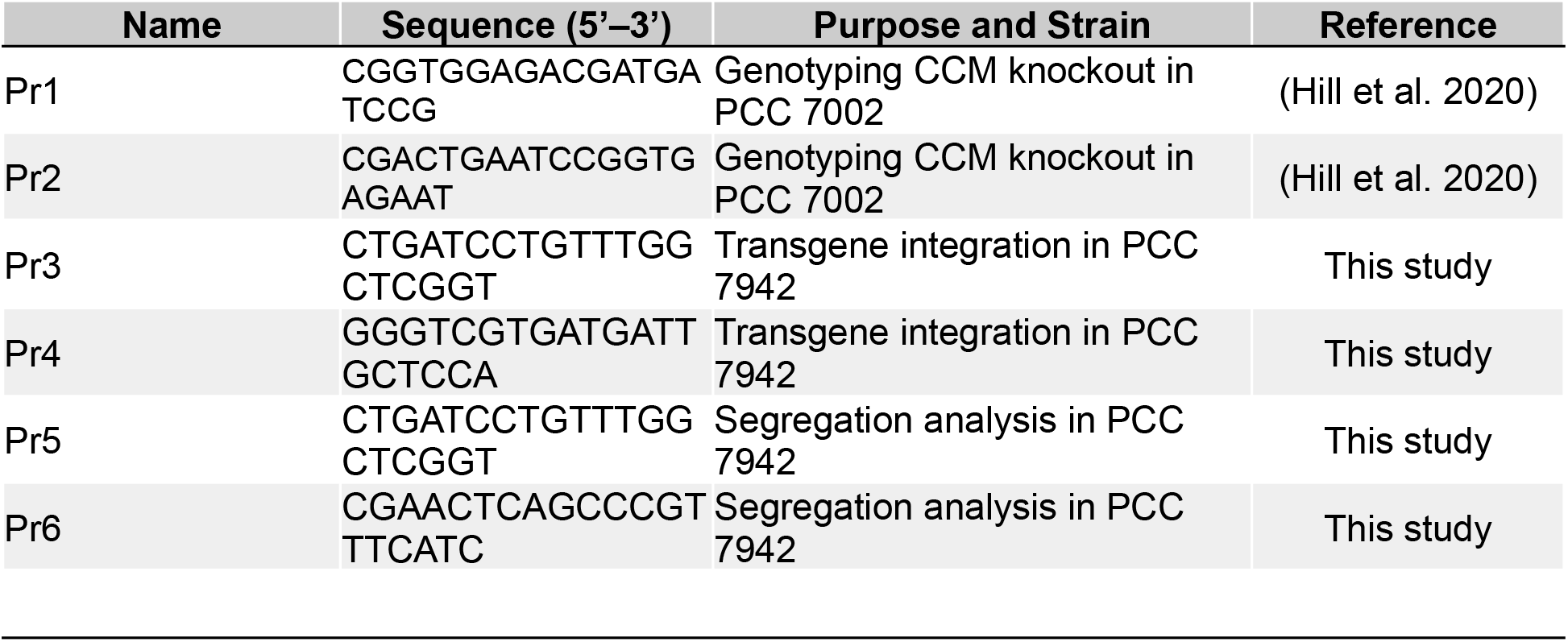
Primers used in this study.

The Δ*ccm*-*7942* strain was constructed similarly to Δ*ccm-1* (PCC 7002), where the region correspondent to genes ccmK to ccmO, known as the *ccm* operon (*ccmKLMNO*-PCC *7942*) was replaced with a spectinomycin resistance cassette using targeted gene deletion via homologous recombination, similar to (Cameron et al. 2013; Clerico et al. 2007). Transformed cells were plated on solid media containing spectinomycin and assessed for transgene segregation by PCR using the primers **Pr5** and **Pr6 (Table 4)**. The genomic DNA was extracted and sequencing confirmed lack of the *ccm* operon.

All PCC 7002 cells were maintained in a modified Guillard’s F/2 medium (Guillard 1975) (F/2 hereafter) supplemented with 5.94 mM NaNO_3_ and 0.25 mM KH_2_PO_4_ (final concentrations). PCC 7942 cells were maintained in Bg-11 medium buffered at pH 7.8 (Allen and Stanier 1968). Strains were maintained in flasks under constant illumination (60 μmol photons m^-2^ s^-1^ white light, spectra provided in **Sup Fig. S9**) with shaking (120 rpm) at 3% CO_2_ and 34 °C. Single colony isolates were stored in F/2 supplemented with 10% dimethyl sulfoxide (D-360-100, Gold Biotechnology, St. Louis, MO, USA) at −80 °C. Cells from the same colony isolate were revived and recultured monthly to avoid mutations following multiple generations of reculturing.

### N Stepdowns (Solid Media)

25 mL cultures were grown in flasks at 3% CO_2_ in N-replete media until absorbance at λ = 750 nm (A_750_) was approximately 0.8. Cells were harvested via centrifugation (5000 x g for 10 minutes) and resuspended in a N-free F/2 medium (where NaNO_3_ was replaced with equimolar NaCl) at final densities of A_750_ = 7 and A_750_ = 21. Cultures were plated in 10 μL volumes on F/2 medium supplemented with 1% Agar and 10 mM Sodium Thiosulfate. Images of plates were captured after 24 hours.

### N Stepdowns (Liquid Media)

Cultures were grown in flasks at 3% CO_2_ in N-replete media until A_750_ was between 0.3 and 0.8. Cells were harvested via centrifugation (5000 x g for 10 minutes) and resuspended in N-deplete F/2 medium (where NaNO_3_ was replaced with equimolar NaCl) or standard F/2 medium (N-replete) at a final density of A_750_ = 0.35, as in Ludwig and Bryant (2012). A_750_ = 0.35 corresponds to 1.25×10^6^ cells/mL on average in these conditions (**Sup Fig**. **S10**). Following resuspension at A_750_ = 0.35, cells were immediately placed at 60 μE white light with shaking (120 rpm), in 1.5 mL volumes in 12-well plates. Gas mixtures (**Table 1**) were produced using a Gas Mixing System (GMS 150, photon systems instruments, Drásov, Czech Republic) and pumped into the headspace at a rate of 0.8 L min^-1^ after bubbling through water.

### Hemacytometry

Images were taken on a Neubauer Grid (Hausser Scientific) using 10 μL of undiluted culture under 20x magnification. A representative image is provided in **Sup Fig. S11**. Cell counts were collected from images using a macro in ImageJ.

### Automated Cell Counting (ImageJ)

Images of cyanobacterial cells were segmented to contain individual 1 mm × 1 mm grids and processed using the following steps in Fiji: (i) background subtraction using the “Subtract Background” function (rolling ball radius = 10 pixels; light background; separate processing), (ii) conversion to a binary mask (“Convert to Mask”), and (iii) particle quantification using “Analyze Particles” with a size threshold of 5–600 pixels^2^. For validation, the macro was compared to a manual cell count shown in **Sup Fig. S11**.

Automated cell counts were used to normalize calculated PBS degradation per cell in **Fig. 1** and **Fig. 2**. Using data generated in **Fig. 2**, we confirmed that A_750_ is a robust predictor of cell number across all genotypes, CO_2_ conditions, and N conditions tested (**Sup Fig. S12**). Therefore, A_750_ was used to normalize calculated PBS degradation in all other experiments.

The number of cell divisions in a given experiment was calculated using the following equation:

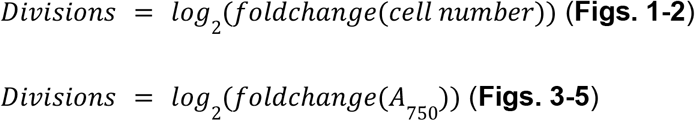

Where *Divisions* reflects the calculated number of divisions after a transition, and *foldchange* (*cell number*) represents the final cell number divided by the initial. Cell number was calculated using hemacytometry, and *A*_*750*_ is the absorbance at wavelength 750 nm.

### Spectrophotometric approximation of PBS

PBS content was approximated using spectrophotometric measurements of live cells according to (Collier and Grossman 1992). Absorbance at 620 nm (corresponding to PBS), and 750 nm (corresponding to cell density) was recorded for live cells. Cells were then heated at 75°C for 8 minutes, and absorbance was re-measured at each wavelength. Approximate PBS content was calculated using the following equation:

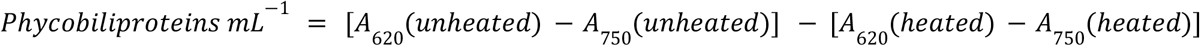

Where *A*_*λ*_ represents absorbance at wavelength λ, and (*unheated*) and (*heated*) represent the cells before and after heating at 75°C, respectively.

### Enzyme kinetic modeling

Enzyme reaction rates were modeled using the Michaelis-Menten equation, assuming no variation of the enzyme concentrations between genotypes. For PEPC, carboxylation rate was calculated as:

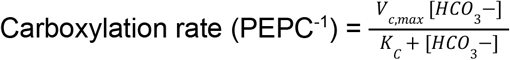

Where *V*_*c,max*_ represents the maximal carboxylation rate of PEPC and *K*_*C*_ represents the catalytic constant for HCO_3_^-^. The constants used for this calculation are provided in **Table 5**.

**Table 5.** PEPC kinetic constants. Values from (Durall et al. 2020).

| Parameter | Value |
| --- | --- |
| Maximal Carboxylation Rate | 14.4 $\mu\text{mol min}^{-1} \text{mg}^{-1}$ |
| Catalytic Constant ($\text{HCO}_3^-$) ($K_c$) | 970 $\mu\text{M}$ |

Bicarbonate concentration was calculated based on the proportion of CO_2_ to HCO_3_^-^ expected at equilibrium in pH 7.8 media (1:50). The concentration of dissolved CO_2_ was calculated using Henry’s Law, according to the following equation:

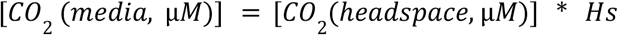

Where *Hs* denotes the Henry’s Solubility constant, (3.5 x 10^-7^ mol kg^-1^ Pa^-1^) for CO_2_.

For Rubisco, a Michaelis-Menten model accounting for two competing substrates (CO_2_ and O_2_) was used and carboxylation and oxygenation rates were calculated as:

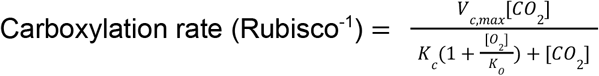

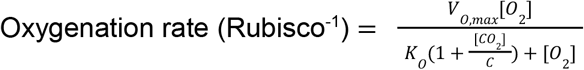

Where *V*_*C,max*_ and *V*_*O,max*_ are the maximal carboxylation and oxygenation rates of Rubisco, and *K*_*C*_ and *K*_*O*_ are the catalytic constants for CO_2_ and O_2_, respectively. The constants used for this calculation are provided in **Table 6**.

**Table 6.**
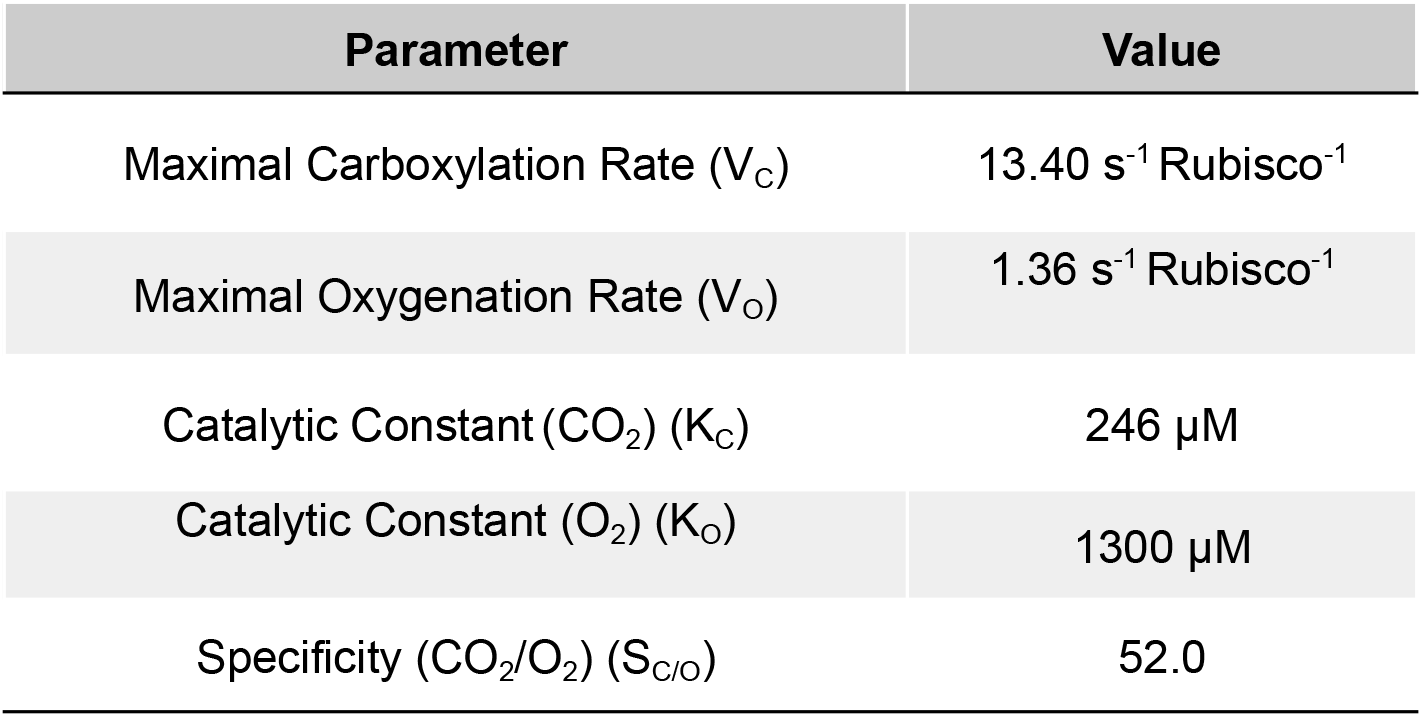
Rubisco (form IB) kinetic constants. Values from (Cummins et al. 2018)

### Statistical analysis and data visualization

All statistical tests, including Student’s T-Tests, ANOVA, and linear regressions were performed in R version 4.0.2. Linear regressions comparing PBS degradation and Rubisco carboxylation rate, oxygenation rate, and PEPC carboxylation rate were performed separately, using the lm() function in R. All graphs were made using the packages *ggplot, ggh4x, cowplot, rvg*, and *officer*.

## Author Contributions

Conceptualization. EVS, SYR, AB

Investigation. EVS, JAG, JA, DT

Resources. JA, PMS, AB, SYR

Funding Acquisition. SYR, AB

Supervision. SYR, AB

Visualization. EVS

Writing - Original draft. EVS

Writing - Review and Editing. EVS, SYR, AB

## Acknowledgements

We thank Jeffrey Cameron for the generous donation of all strains in the PCC 7002 genetic background, and for his helpful and thoughtful feedback on this study. We thank Professors Arthur Grossman, Devaki Bhaya, Daniel Ducat, Berkley Walker, and James Moran for helpful discussions on this project. We further thank the Rhee, Burlacot, Grossman, and Bhaya labs for valuable discussions on experimental design and the writing of the manuscript. This work was conducted in part on the ancestral land of the Muwekma Ohlone Tribe which was and continues to be of great importance to the Ohlone people, and on the ancestral, traditional, and contemporary lands of the Anishinaabeg – Three Fires Confederacy of Ojibwe, Odawa, and Potawatomi peoples. The work was also conducted at Michigan State University that occupies the ancestral, traditional, and contemporary Lands of the Anishinaabeg – the Three Fires Confederacy of Ojibwe, Odawa, and Potawatomi peoples.

## Funding

This work was supported in part by Carnegie Science (to AB), the US NIH’s NIGMS Cellular and Molecular Biology Training Program (grant T32GM007276), the U.S. National Science Foundation grants (IOS-2312181, IOS-2406533, IOS-1546838, MCB-1617020, MCB-2052590, MCB-1916797, MCB-2420360, OISE-2434687, DBI-2213983 to SYR) and U.S. Department of Energy, Office of Science, Office of Biological and Environmental Research, Genomic Science Program grants (DE-SC0018277, DE-SC0023160, DE-SC0008769, and DE-SC0021286 to SYR and DE-SC0019417 to AB).

## Conflicts of Interest

The authors declare no conflicts of interest

## Data Availability

The data from this article will be shared upon reasonable request to the corresponding authors.

## Notes

### Competing Interest Statement

The authors have declared no competing interest.

