## Supplemental Figures for "CO_2_ fixation mediated by the carbon concentrating mechanism enables a rapid response to nitrogen deprivation in cyanobacteria"

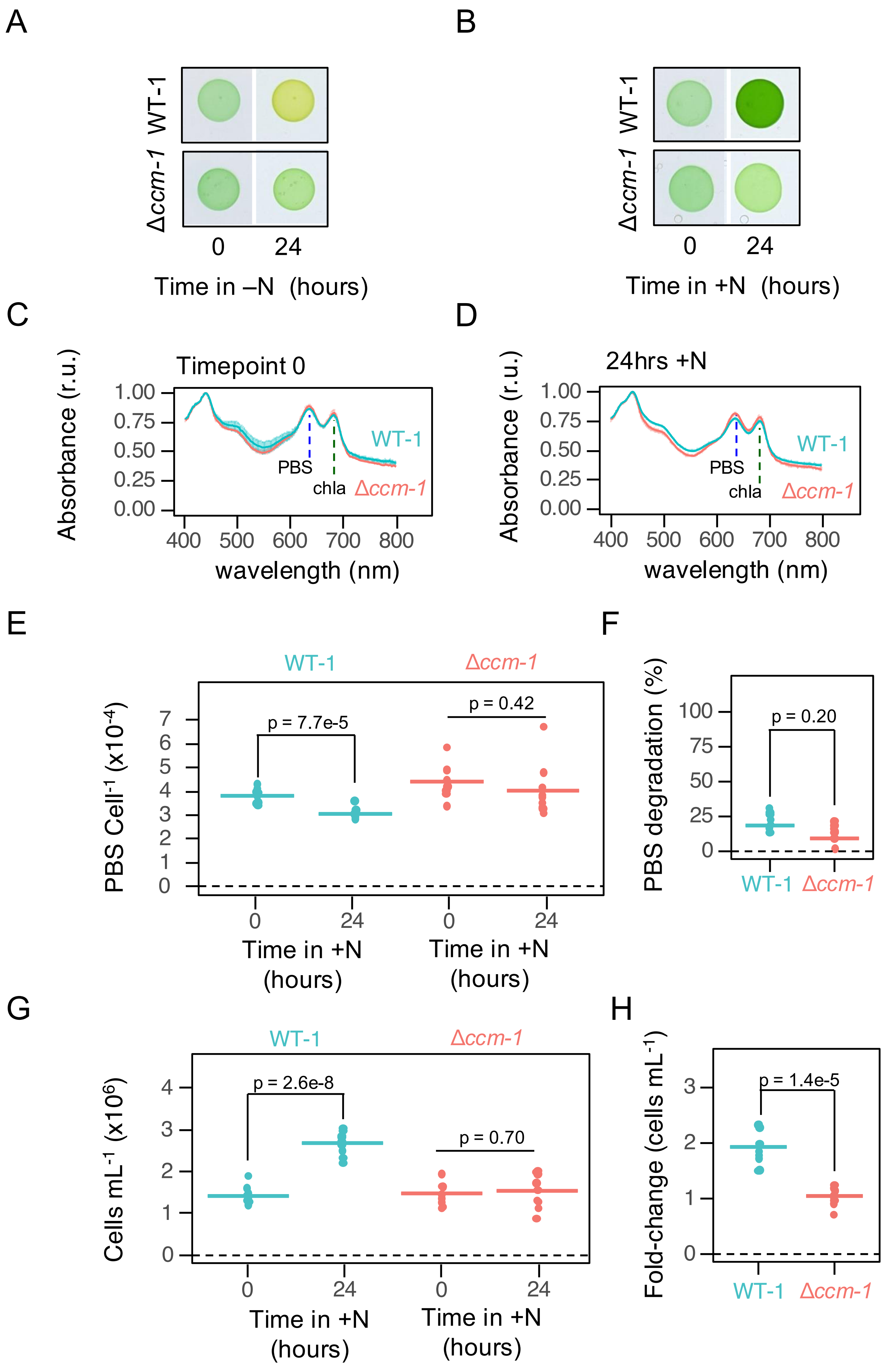
A
WT-1
Δccm-1
0
24
Time in –N (hours)
B
WT-1
Δccm-1
0
24
Time in +N (hours)
C
Timepoint 0
Absorbance (r.u.)
1.00
0.75
0.50
0.25
0.00
400
500
600
700
800
wavelength (nm)
WT-1
PBS
chla
Δccm-1
D
24hrs +N
Absorbance (r.u.)
wavelength (nm)
WT-1
PBS
chla
Δccm-1
E
WT-1
Δccm-1
PBS Cell$^{-1}$ (x10$^{-4}$)
p = 7.7e-5
p = 0.42
Time in +N (hours)
F
PBS degradation (%)
p = 0.20
WT-1
Δccm-1
G
WT-1
Δccm-1
Cells mL$^{-1}$ (x10$^{6}$)
p = 2.6e-8
p = 0.70
Time in +N (hours)
H
Fold-change (cells mL$^{-1}$)
p = 1.4e-5
WT-1
Δccm-1

**Supplemental Figure S1. Physiological changes caused by the transition from HC to LC.** A-B) Representative images of WT-1 and Δ*ccm-1* cells taken before and after 24 hours without N (A) and with N (B) on solid media at 0.04% $CO_2$. Panel A is reproduced from **Fig. 1B** to facilitate comparison with panel B. C-D) Absorbance spectra for WT-1 and Δ*ccm-1* cells before (C) and 24 hours after (D) transfer to 0.04% $CO_2$ (LC). Absorbance values were rescaled so that max absorbance = 1. Because this normalization was applied, absorbance values are given as relative units (r.u.). Two vertical dashed lines show representative wavelengths used to estimate the abundance of phycobilisomes (PBS) and chlorophyll A (chla). Lines represent means of 3 independent biological replicates. E-F) Changes in per-cell PBS content over 24 hours in liquid culture at LC. Per-cell PBS content is shown at timepoint 0 and 24 hours in C. The decrease in per-cell PBS content (PBS degradation) after 24 hours is shown in F. G-H) Changes in cell number after 24 hours at LC. Cell densities are shown at timepoint 0 and 24 hours in G. The change in cell density over 24 hours is shown in H. In E-H, each dot represents a single biological replicate, and horizontal bars represent the mean. *N = 3* independent experiments were performed (*n = 2-3* biological replicates per genotype per experiment). p-values were calculated using a Student's t-test.

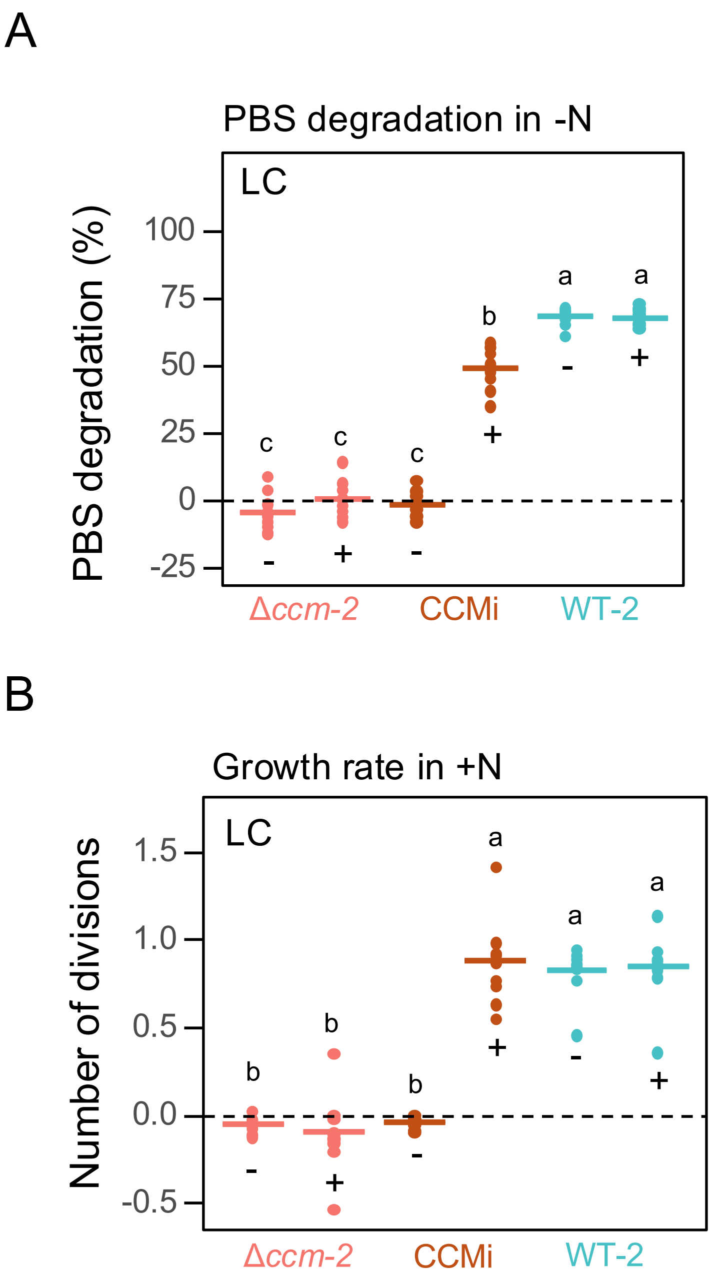


**Supplemental Figure S2. Genetic complementation rescues PBS degradation in CCM mutants.** A) Phycobilisome (PBS) degradation after 24 hours of nitrogen (N) deprivation in WT-2, Δ*ccm-2*, and CCMi strains at LC (0.04% $CO_2$) with (+) and without (-) Isopropyl-β-D-galactopyranoside (IPTG). B) Number of cell divisions after 24 hours in N-replete media for the same experiment shown in A. Each data point in A and B represents a single biological replicate from $N = 3$ independent experiments ($n = 3$ biological replicates per genotype per experiment per condition). The horizontal line represents the mean of all biological replicates. Letters represent significance codes from a two-way ANOVA and Tukey's Honest Significance Difference (HSD) test. Different letters indicate a statistically significant difference at p-value < 0.05.

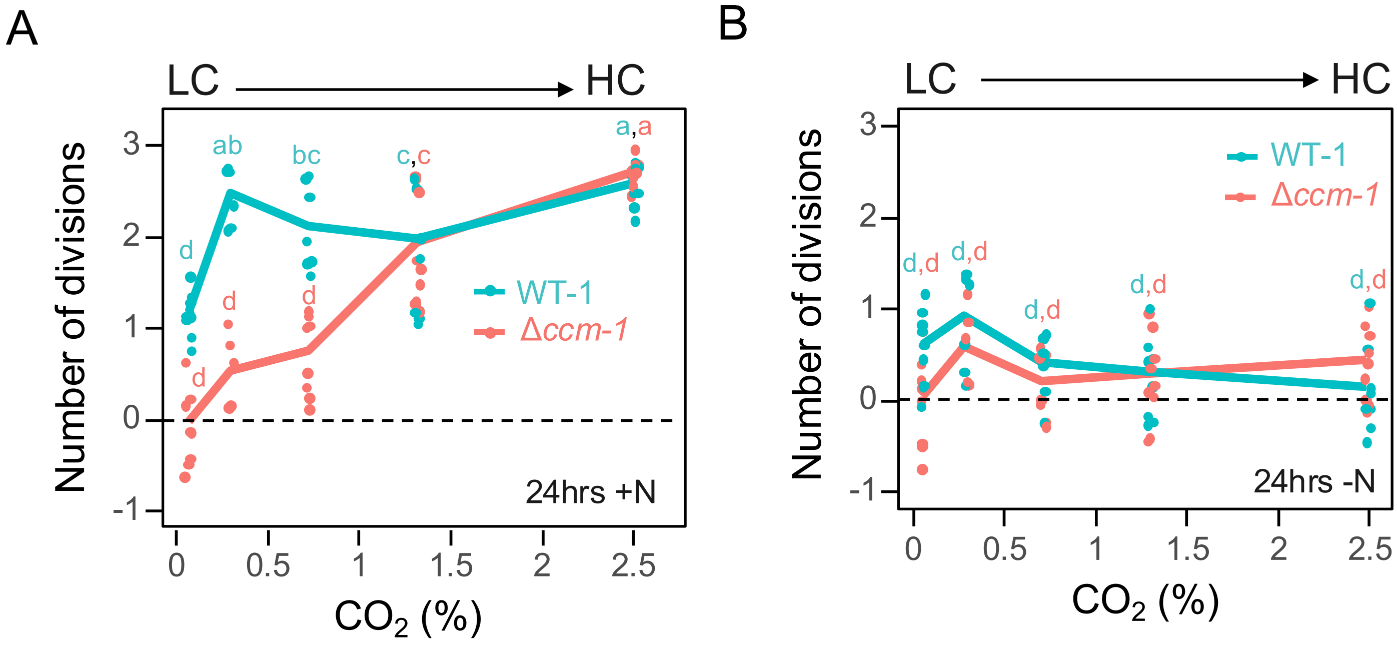


**Supplemental Figure S3. Cell growth across $CO_2$ conditions.** A) Number of cell divisions after 24 hours in nitrogen (N)-replete media for WT-1 and Δ*ccm-1* cells across various $CO_2$ conditions. Data in (A) were collected from the samples shown in **Fig. 2.** B) Number of cell divisions occurring after 24 hours in N-deplete media for WT-1 and Δ*ccm-1* cells across various $CO_2$ conditions. The $CO_2$ conditions shown are 0.04% (LC), 0.23%, 0.7%, 1.3%, and 2.5% (HC). Data in (A) were collected from cells from the same experiment shown in panel B, which were transitioned to N-replete medium instead of N-deplete medium. Each data point represents a single biological replicate. Data are from *N = 3* independent experiments (*n = 3* biological replicates per genotype per condition per experiment). The line indicates the mean of all biological replicates. Letters represent significance codes from a three-way ANOVA and Tukey's Honest Significance Difference (HSD) test. Different letters indicate a statistically significant difference ($p < 0.05$).

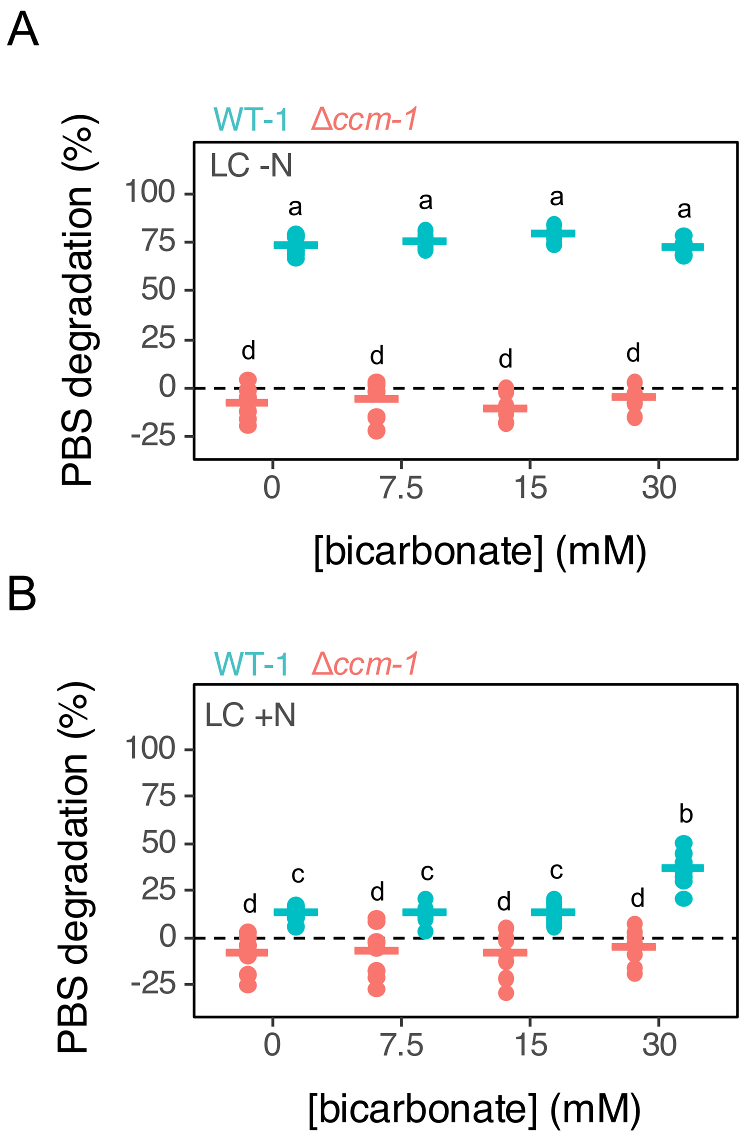


**Supplemental Figure S4. PBS degradation in CCM mutants is not bicarbonate-dependent.** Normalized PBS degradation 24 hours of nitrogen (N) deprivation (A) or in N-replete media (B) for WT-1 and Δccm-1 cyanobacteria at LC supplemented with bicarbonate ranging from 0–30 mM. Data in A and B are from ($N$ = $3$ independent experiments, each with $n$ = $3$ biological replicates per genotype per condition). In A and B, each data point represents a biological replicate and horizontal bars indicate the mean of all biological replicates for each condition and strain. Letters represent significance codes from a three-way ANOVA and Tukey's Honest Significance Difference (HSD) test performed across (A) and (B). Different letters indicate a statistically significant difference at p-value < 0.05.

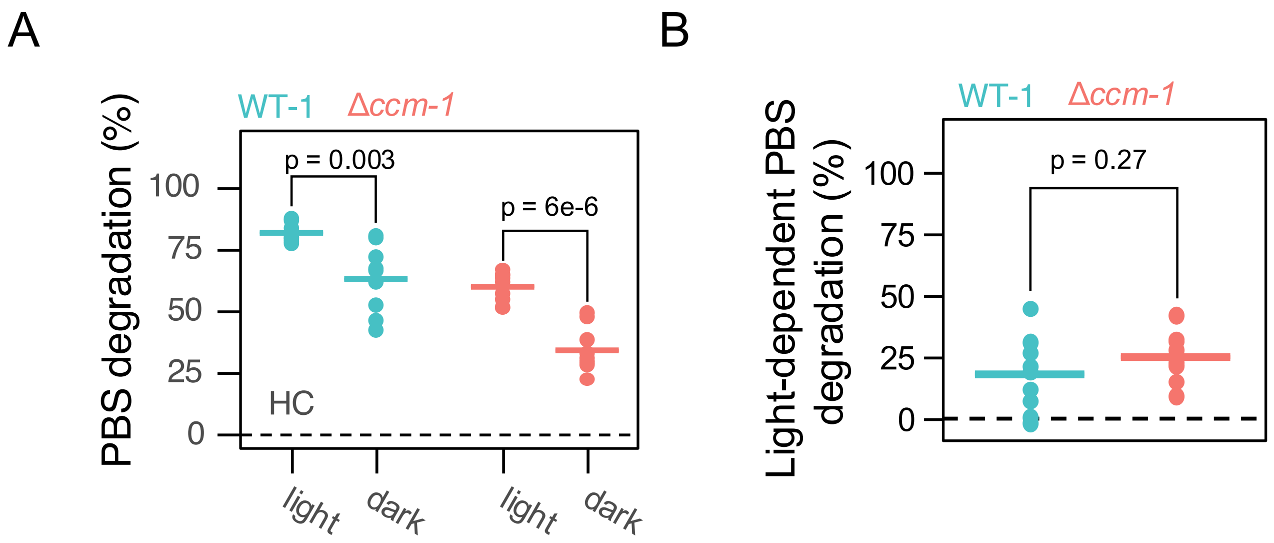


**Supplemental Figure S5. Light energy-dependent bleaching for WT-1 and Δ*ccm-1*.** (A) Normalized PBS degradation following 24 hours of N deprivation in WT-1 and Δ*ccm-1* under 60 μE or 0 μE of white light (light and dark, respectively). (B) Light-dependent PBS degradation (difference between light and dark, from A) is shown for WT-1 and Δ*ccm-1*. Each data point represents a single biological replicate. Data in A and B are from *N = 3* independent experiments (*n = 3* biological replicates per experiment per genotype per condition). p-values were calculated using a Student's t-test.

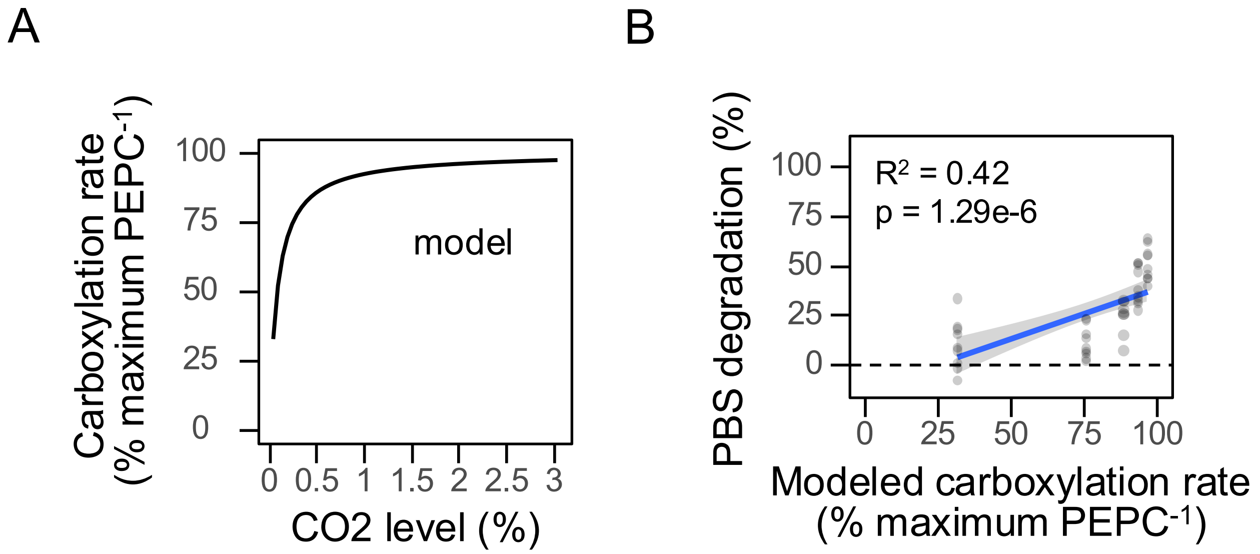


**Supplemental Figure S6. Phosphoenolpyruvate (PEP) carboxylase activity correlates with PBS degradation in CCM mutants.** A) Modeled carboxylation rates per PEP carboxylase (PEPC) enzyme using kinetic parameters specific to PCC 7002 PEPC (**Methods**). B) Comparison of PBS degradation in Δccm-1 mutant cells (from **Fig. 2**) to predicted PEPC carboxylation rates. Data from Fig. 2 are overlaid in light grey. Adjusted $R^2$ values are shown for a linear regression comparing PBS degradation (%) with modeled PEPC carboxylation rates. The significance (p) of the F-statistic is listed.

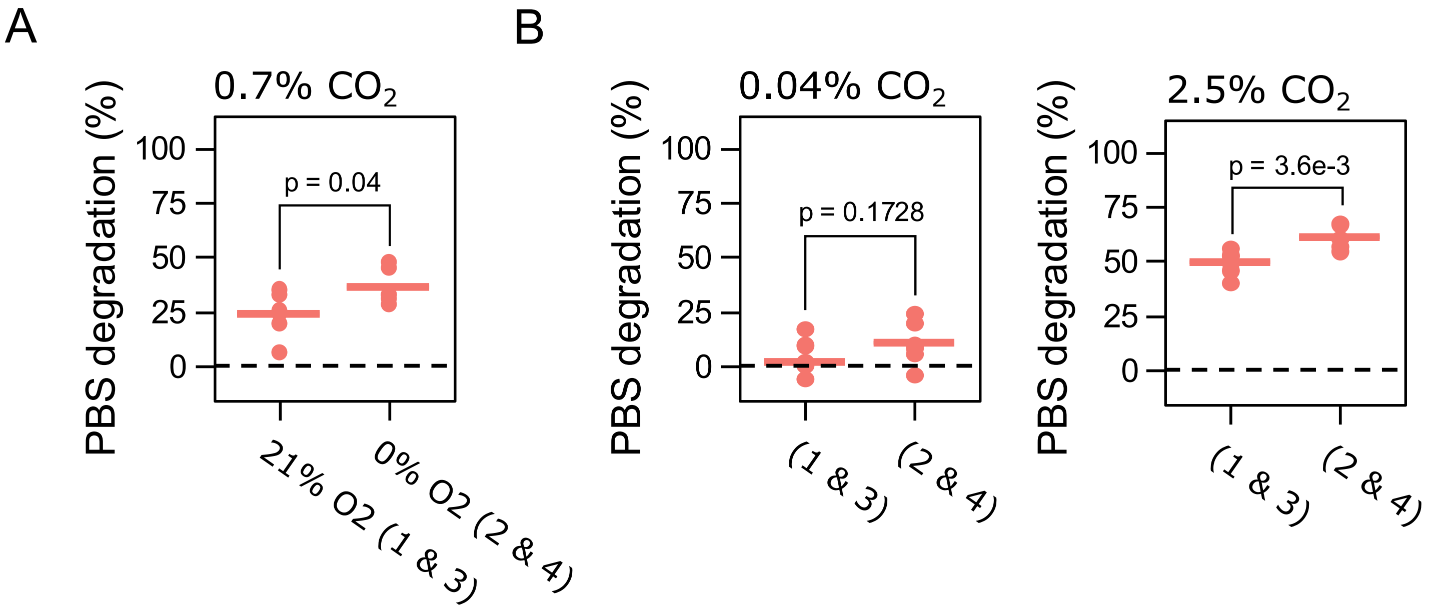


**Supplemental Figure S7. Non-normalized data corresponding to Figure 3D.** A) PBS degradation of Δ*ccm-1* cells after 24 hours of N deprivation for Δ*ccm-1* cells at 0.7% $CO_2$ with and without $O_2$. The data was compiled from 4 experiments on 4 days, with 21% $O_2$ tested on days 1 and 3, and 0% $O_2$ tested on days 2 and 4. B) PBS degradation of Δ*ccm-1* cells from the same experiments after 24 hours of N deprivation at 0.04% $CO_2$ (left) and 2.5% $CO_2$ (right). Each data point represents a single biological replicate (*n = 3* biological replicates per genotype per condition per experiment, *N = 2* experiments). The horizontal line represents the mean of 6 biological replicates. The p-value from Student's t-test is shown (p).

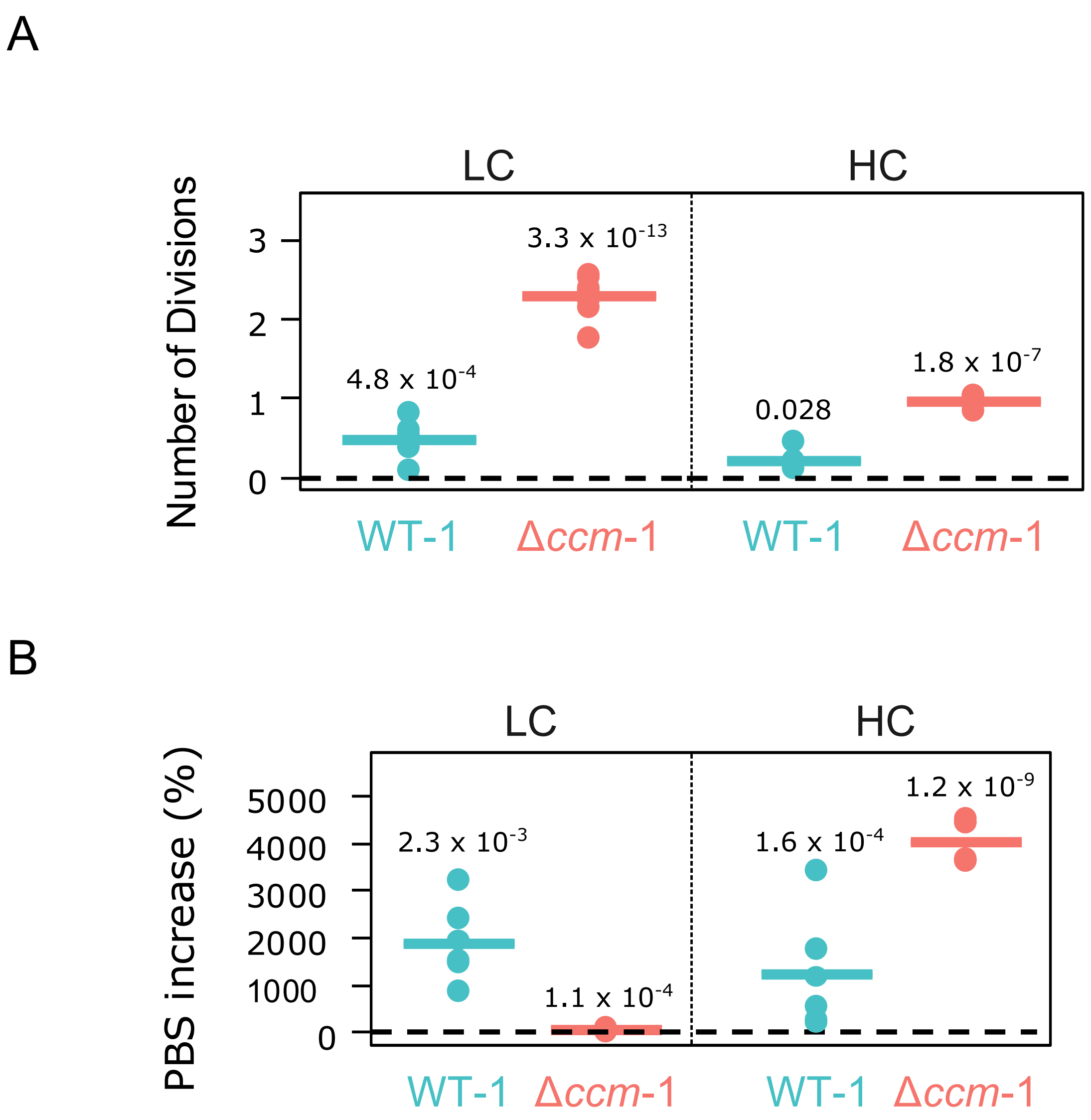


**Supplemental Figure S8. Recovery of WT-1 and Δ*ccm-1* from N deprivation.** Growth (A) and PBS synthesis (B) from cells transitioned to N-replete media at HC after 48 hours of N deprivation at LC (left) or HC (right). Recovery is represented as A) number of divisions after 36 hours of N resupply and B) increase in PBS per cell after 36 hours of N resupply. Each data point represents a single biological replicate. Solid lines indicate the mean of all biological replicates from $N = 2$ independent experiments each with $n = 3$ biological replicates per genotype. Numbers represent p-values from a one-sample Student's t-test comparing the mean to 0. Data were collected from the same cultures in **Fig. 3**, after the 48 hour N deprivation was completed.

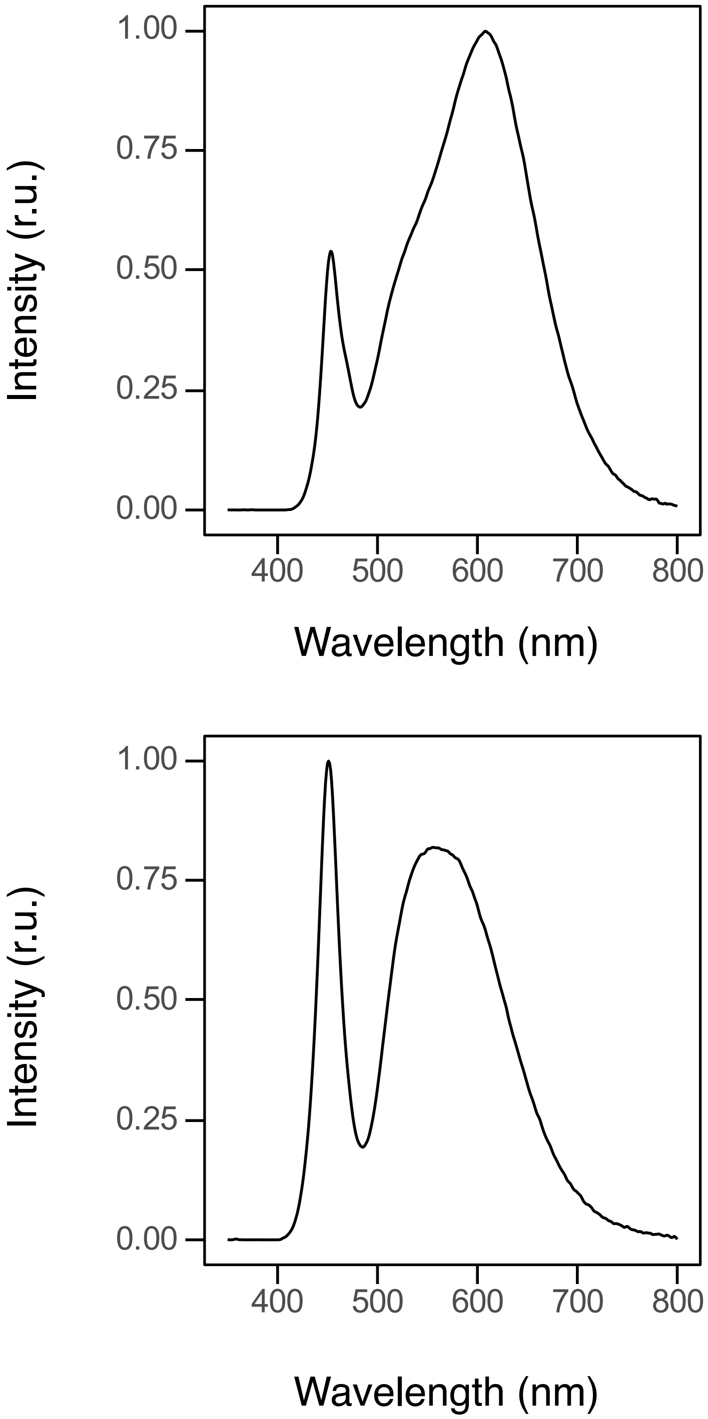


**Supplemental Figure S9. Light spectra used throughout this study.** (A) Light spectrum of the growth chamber used for preculturing cells for all experiments. This growth chamber was also used for experiments in **Figs. 3, Sup Fig. 8** (B) Light spectrum of the LED panels used for experiments in **Figs. 1, 2, 4, 5, Sup Figs. S1, S2, S3, S4, S5, S6, S7.** r.u. = relative units.

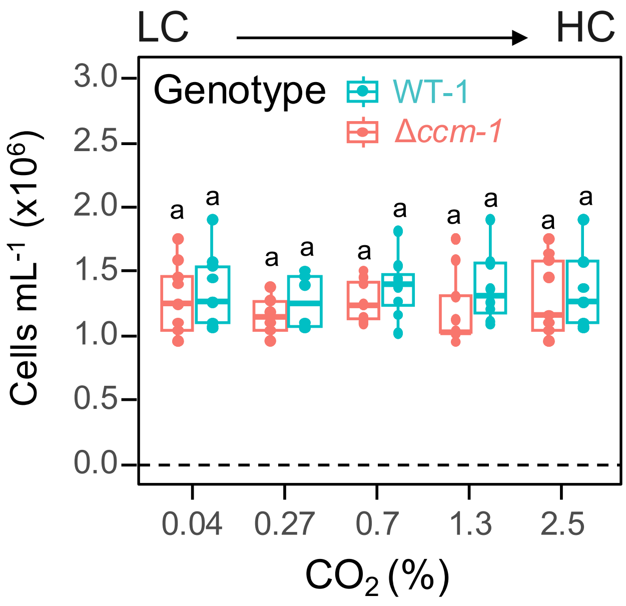


**Supplemental Figure S10. Starting cell densities for N deprivation experiments.** Starting cell densities from the experiments shown in **Fig. 2.** Densities were measured by hemocytometry from cultures after dilution to a starting $A_{750}$ = 0.35. Box-and-whisker plots show the median (center line), first and third quartiles (box limits), and whiskers extending to 1.5× the interquartile range. Individual data points are overlaid. Each dot represents a single starting culture from *N = 3* independent experiments (*n = 3* replicates per genotype per condition per experiment). Letters represent significance codes from a two-way ANOVA and Tukey's Honest Significance Difference (HSD) test. Different letters indicate a statistically significant difference at p-value < 0.05.

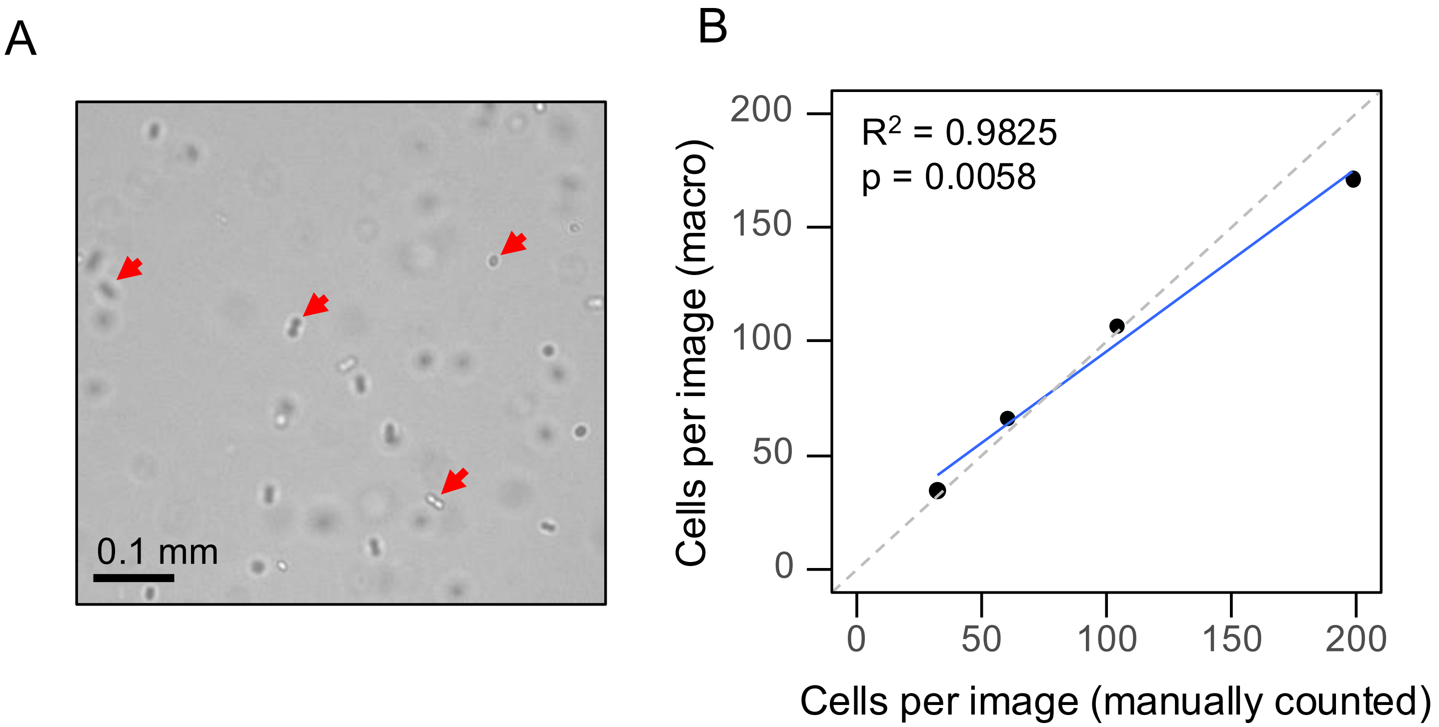


**Supplemental Figure S11. Performance of automated hemocytometry macro.** A) Representative image of cyanobacterial cells in a Neubauer chamber. Arrowheads indicate cells ranging from completely out of focus (top left) to in focus (bottom right). B) Cells per image calculated using the macro described in methods (y-axis) compared with manual count obtained by scrolling through all focal planes (x-axis). The blue line represents a linear regression; adjusted $R^2$ and the p-value of the fit are shown (p). Each data point represents the mean of 5 independent manual and automated measurements. The diagonal dashed line has a slope of 1 and intersects at y = 0.

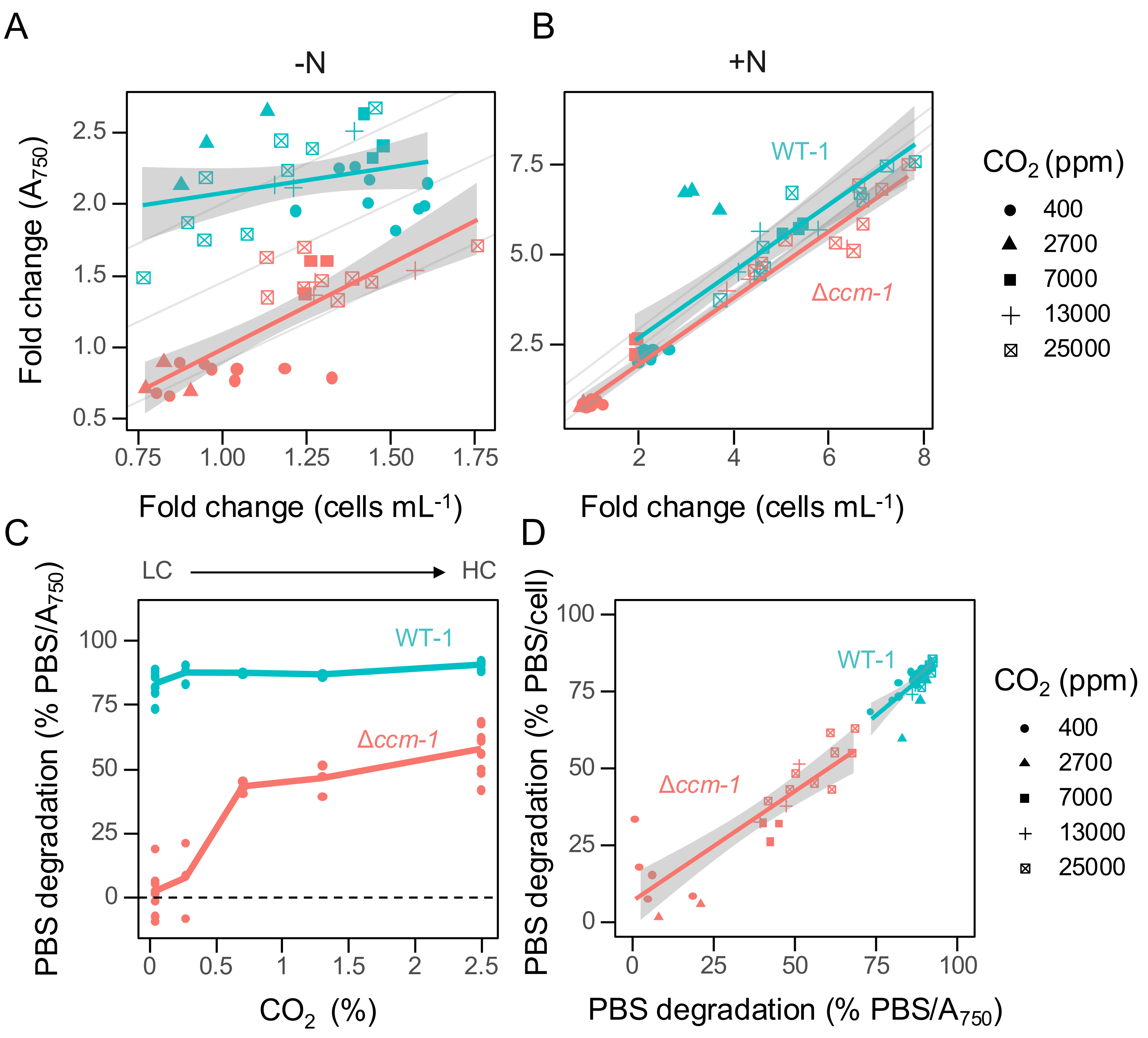


**Supplemental Figure S12. Relationship between cell number and $A_{750}$ across strains and conditions** A-B) Comparison of measured growth (fold change in cells $mL^{-1}$) and estimated growth (fold change in Absorbance at 750 nm [A750]) for WT and Δ*ccm-1* cells following 24 hours of growth in nitrogen (N)-deplete (A) or N-replete (B) media under various $CO_2$ conditions. A linear regression (solid line) is shown for each genotype in both N conditions. The shaded area represents the 95% confidence interval. C) Normalized phycobilisome (PBS) degradation following 24 hours of N deprivation for WT and CCM mutant cyanobacteria across various $CO_2$ conditions. The $CO_2$ conditions shown are 0.04% (LC), 0.23%, 0.7%, 1.3%, and 2.5% (HC). PBS degradation was normalized by $A_{750}$. D) Comparison of PBS degradation normalized by A750 (as in panel C) versus PBS degradation normalized by cell number (as in **Fig. 2**).

**Supplemental Figure S12 continued.** A linear regression (solid line) is shown for each genotype. The shaded area represents the 95% confidence interval. Data in A-D were derived from the same experiment as **Fig. 2**.
